# Single-nucleus multiomic analysis reveals modulation of the gene regulatory circuit landscape in pituitary cell types during mouse estrous cycle

**DOI:** 10.64898/2026.06.16.732671

**Authors:** Zidong Zhang, Wan Sze Cheng, Yangfan Jin, Luisina Ongaro, Gregory R. Smith, Hanna Pincas, Natalia Mendelev, Galia Strupinsky, Carlos Agustin Isidro Alonso, Xiang Zhou, Emilie Brûlé, Michel Zamojski, Judith L. Turgeon, Elena Zaslavsky, Daniel J. Bernard, Frederique Ruf-Zamojski, Stuart C. Sealfon

**Author notes:** Authors contributed equally to this work (Zidong Zhang, Wan Sze Cheng).

## Abstract

The estrous cycle transcriptional and chromatin dynamics in pituitary cell types have not been investigated. We report single-nucleus multiomics assays in 18 adult mouse pituitaries (102,069 post quality-control nuclei) across 6 cycle time points. Differential analysis revealed cycle stage-dependent epigenetic and transcriptional remodeling across the major pituitary cell populations. In gonadotropes and lactotropes, we identified temporal patterns of differential gene expression linked to various biological processes, notably neuronal and synaptic-related ontologies. Pseudotime trajectory analysis was consistent with a rapid transition of individual gonadotropes through different cellular states. In gonadotropes and lactotropes, we uncovered gene regulatory circuits whose activity varies between consecutive cycle time points. We experimentally validated a gonadotrope ETS2-driven circuit that differentially regulates *Fshb* gene expression between 2 and 9 am on estrus. Our data and analyses, available at https://rstudio-connect.hpc.mssm.edu/snpit_estrous_browser/, provide a window into gene regulatory mechanisms underlying estrous cycle stage transitions.

**Highlights:**

- Pituitary cells show chromatin and transcriptome plasticity across the estrous cycle
- We identify stage-modulated gene regulatory circuits across pituitary cell types
- Gonadotrope DEGs form distinct pathway-annotated temporal trajectory clusters
- We validate a gonadotrope ETS2-driven circuit regulating the *Fshb* gene

**eTOC blurb:** Zhang *et al.* conduct a single-nucleus multiomics analysis of mouse pituitaries *in vivo* across the estrous cycle. They reveal an epigenetic and transcriptomic plasticity in pituitary cell populations. In gonadotropes and lactotropes, they demonstrate that DEGs clustered by temporal trajectories are enriched for distinct biological processes and identify stage-modulated gene regulatory circuits. They experimentally validate a gonadotrope ETS2-driven circuit regulating *Fshb* expression. Their dynamic molecular atlas of the cycling pituitary captures key *cis*-regulatory mechanisms underlying estrous cycle stage transitions.

## Introduction

The reproductive cycle is essential for species survival, as it determines the timing of female fertility. Referred to as the estrous cycle in rodents, it is orchestrated by the hypothalamic-pituitary-gonadal (HPG) axis. The pituitary gland plays a central role in coordinating the cycle through the release of the gonadotropins, luteinizing hormone (LH) and follicle-stimulating hormone (FSH; ^1^). During the estrous cycle, the complex and carefully timed hormonal interplay between the hypothalamus, the pituitary, and the gonads leads to a mid-cycle LH surge and biphasic FSH surges, inducing ovulation and ovarian follicle development, respectively ^2–5^.

Previous *in vivo* and *in vitro* studies in rodents showed that pituitary gonadotrope cells undergo functional and organizational changes in response to GnRH and other stimuli during the estrous cycle, including alterations in the number of GnRH receptors and in the abundance of LH and FSH secretory granules, morphological rearrangements such as cellular migration and the development of cellular projections towards the capillary network, which facilitate vesicle trafficking and secretion (for review, ^6^). The gene regulatory mechanisms underlying the dynamic changes induced throughout the estrous cycle in gonadotropes (GT) and other pituitary cell populations have not been thoroughly investigated. Elucidating the mechanisms that underpin pituitary cell plasticity is essential to determine the molecular basis of the physiological changes associated with the reproductive cycle, and has clinical implications for the treatment of infertility and the potential development of new hormonal contraceptives.

Recently, the application of single-cell (sc) and single-nucleus (sn) transcriptomics has enabled researchers to resolve cellular heterogeneity of the pituitary gland in normal and pathological states or during development, as well as to detect gene expression changes in disease or in response to physiological stimuli (^7–9^ ; for review, see ^10^). Sn multiome assays that simultaneously measure the transcriptome and chromatin accessibility within the same individual cells provide deep insight into the molecular state of distinct cells and cell types, and allow the characterization of cell type-specific gene regulatory mechanisms ^11–15^. When combined with recently developed bioinformatic methods, sc multiomics assays can resolve the gene regulatory circuit (GRC) triads comprising a gene, its *cis*-regulatory transcription factor binding locus, and the regulatory transcription factor, that together orchestrate gene expression dynamics ^16,17^.

The estrous cycle is divided into 4 stages (metestrus, diestrus, proestrus, and estrus), identified by vaginal smear cytology and characteristic gonadal hormone levels ^18,19^. Here, to resolve the gene regulatory mechanisms underpinning pituitary cell plasticity throughout the murine estrous cycle, we carried out sn multiomics assays of the transcriptome and chromatin accessibility in pituitary samples obtained throughout the estrous cycle from mice that were categorized via vaginal cytology and hormonal status. This study revealed the temporal dynamics of gene expression, pathway activity, chromatin accessibility, and GRC activity across the estrous cycle. While focusing on gonadotropes, we captured the gene regulatory dynamics across the major pituitary cell types, finding that broad changes occur throughout pituitary cell types.

## Results

### Estrous cycle assignment and data generation

A schematic of the overall study design is shown in **Fig. 1A**. To examine the dynamic transcriptomic and chromatin accessibility landscapes of mouse pituitaries across cell types, we collected the glands at specific stages of the estrous cycle, which were determined based on vaginal cytology and hormonal status evaluation. Mice housed in 14 hours light/10 hours dark with lights on at 4 am were selected at the following six cycle time points: proestrus 9 am, 6 pm, and 11 pm; estrus 2 am and 9 am, and diestrus 9 am. Pituitaries were harvested, flash frozen, and stored at -80 °C until assay. Vaginal cytology evaluated the relative proportions of cell types present in the vaginal smear (leukocytes, cornified epithelial cells, and nucleated epithelial cells; ^19^), as detailed in the Methods. Serum gonadotropin (LH and FSH) levels, along with inhibin Alpha and inhibin Beta levels measured at the time of sacrifice, were used as biomarkers to confirm accurate assignment of the estrous cycle stage or substage (**Fig. 1B** and **Table S1**; for reference, see ^20,21^. For the sn multiome assays, each pituitary was homogenized individually, and its nuclei were isolated and subjected to snRNAseq and snATACseq simultaneously for subsequent QC and analyses. **Table S2** provides the number of pituitaries used per time point and the number of extracted nuclei.

**Figure 1.**
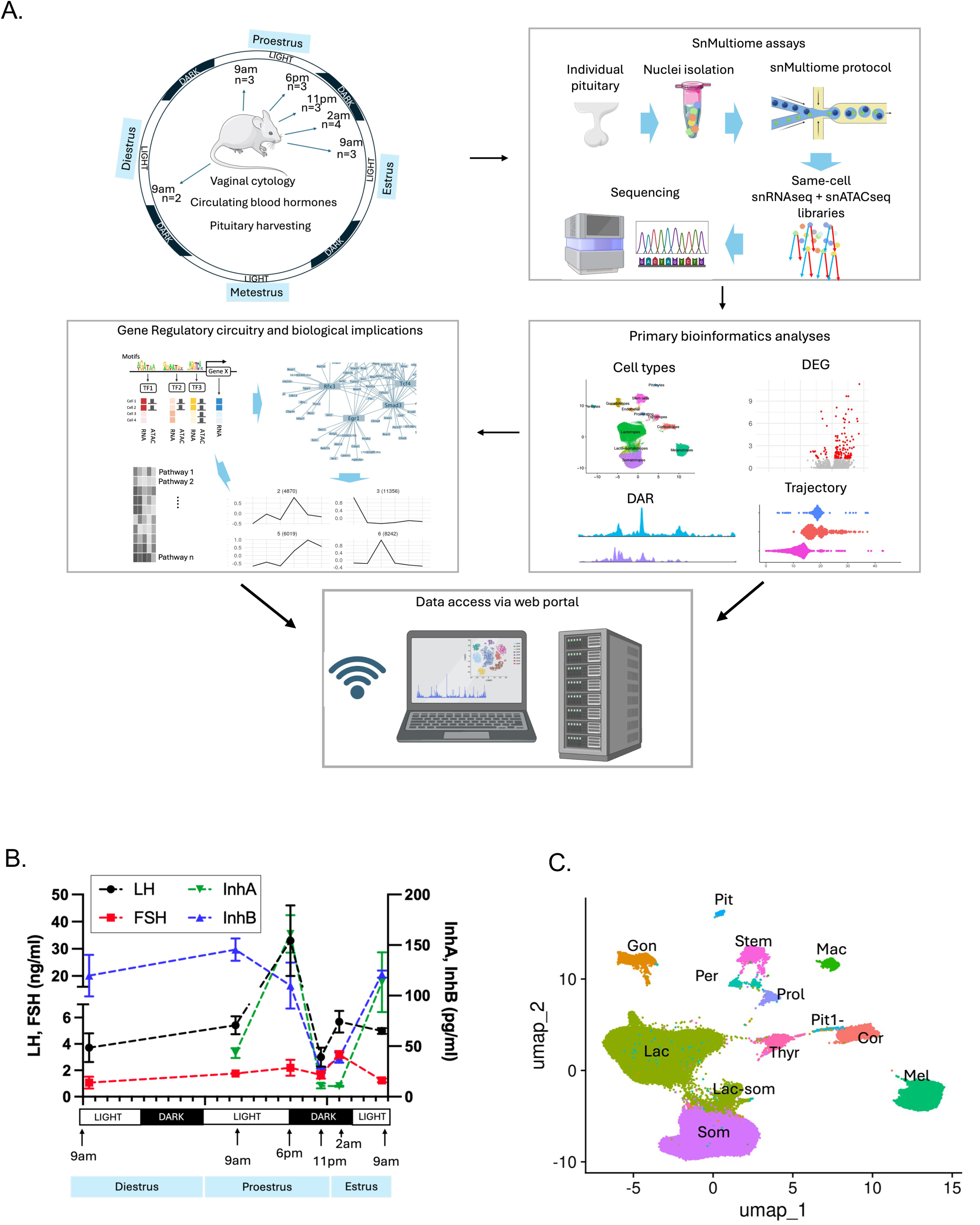
Sn multiomics analysis of the mouse pituitary across the estrous cycle - study design and data overview. **A.** Schematic of the experimental procedure. The major stages depicted are: 1) animal staging and pituitary collection, 2) same-cell scMultiome assay from isolated nuclei, 3) bioinformatics analyses including cell type identification, identification of differentially expressed genes (DEGs), differential chromatin accessibility regions (DARs), and pseudotime trajectory analysis, 4) inference of gene regulatory circuits (GRCs) and their biological implications, 5) access to snRNAseq data, snATACseq data, and integrated datasets via a web portal. **B.** Serum levels of the gonadotropin hormones, Inhibin A, and Inhibin B across the estrous cycle. **C.** Integrated UMAP of the snRNAseq/ATACseq data derived from the 18 samples collected throughout the estrous cycle. Cell type cluster identification is based on established markers for each cell type. Sequencing metrics are provided in **Table S3**.

### Multiomics data processing and identification of pituitary cell types

Data from all samples were evaluated using stringent laboratory-established quality control (QC) standards ^11,14,22^ for both snRNAseq and snATACseq and by visual inspection of Uniform Manifold Approximation and Projection (UMAP) patterns. The data from all 18 samples used for downstream analyses (**Fig. 1A**) passed overall data QC metrics and showed consistent UMAP distributions. These samples included 102,069 cells that passed QC for both assays ^11,14^. Each dataset had more than 5,000 cells analyzed per sample, ∼2,000 genes/cell, a median between 5,600 and 24,000 high-quality ATAC fragments per cell, and transcription start site (TSS) enrichment scores above 9 (**Table S3**). Data were processed using Cell Ranger ARC 1.0.0 (10x Genomics), integrated using the Seurat V4 pipeline, and visualized in UMAP plots by individual and merged data types. To merge data types, the genome-wide chromatin accessibility data were converted to gene-level data, and the converted ATACseq data were integrated with the RNA expression data using nearest neighbor analysis. Cell clusters were annotated using SingleR for label transfer using a reference mouse data atlas ^12^. Both snATACseq and merged UMAPs led to the identification of the same 12 cell populations as the snRNAseq, namely gonadotropes, lactotropes, somatotropes, lacto-somatotropes, corticotropes, thyrotropes, melanotropes, stem cells, proliferating cells, endothelial cells, pituicytes, and pericytes (**Fig. 1C**, **Fig. S1A**), with good correspondence of the pituitary cell type clusters in both assay modalities. No significant changes in pituitary cell type proportions were observed among samples obtained at different estrous cycle time points (**Fig. S2** and **Table S4**).

### Cycle stage-dependent changes in RNA expression and chromatin accessibility across pituitary cell types

We investigated the genome-wide chromatin accessibility and gene expression changes throughout the estrous cycle in the major pituitary cell types. For each cell type, we determined the number of differentially expressed genes (DEGs) and of differentially accessible chromatin regions (DARs) at each of the six estrous cycle stage time points in comparison to the previous time point (**Fig. 2**). Despite gonadotropes being the main pituitary cell type contributing to estrous control, we observed a surprisingly large number of regulatory events in the other pituitary cell types. Across all time points, gonadotropes and lactotropes had the largest number of DEGs, with a total of 3,656 DEG events and 2,143 unique DEGs in the gonadotropes vs. a total of 2,021 DEG events and 1,031 unique DEGs in the lactotropes (**Fig. 2A**). Lactotropes showed the highest number of DARs, with a total of 71,071 DAR events and 39,616 unique loci, followed by somatotropes (27,092 DAR events, 17,659 unique loci) and gonadotropes (1,391 DAR events, 1,274 unique loci; **Fig. 2B**). The full lists of DEGs and DARs per cell type are provided in **Table S5** and **Table S6**, respectively.

**Figure 2.**
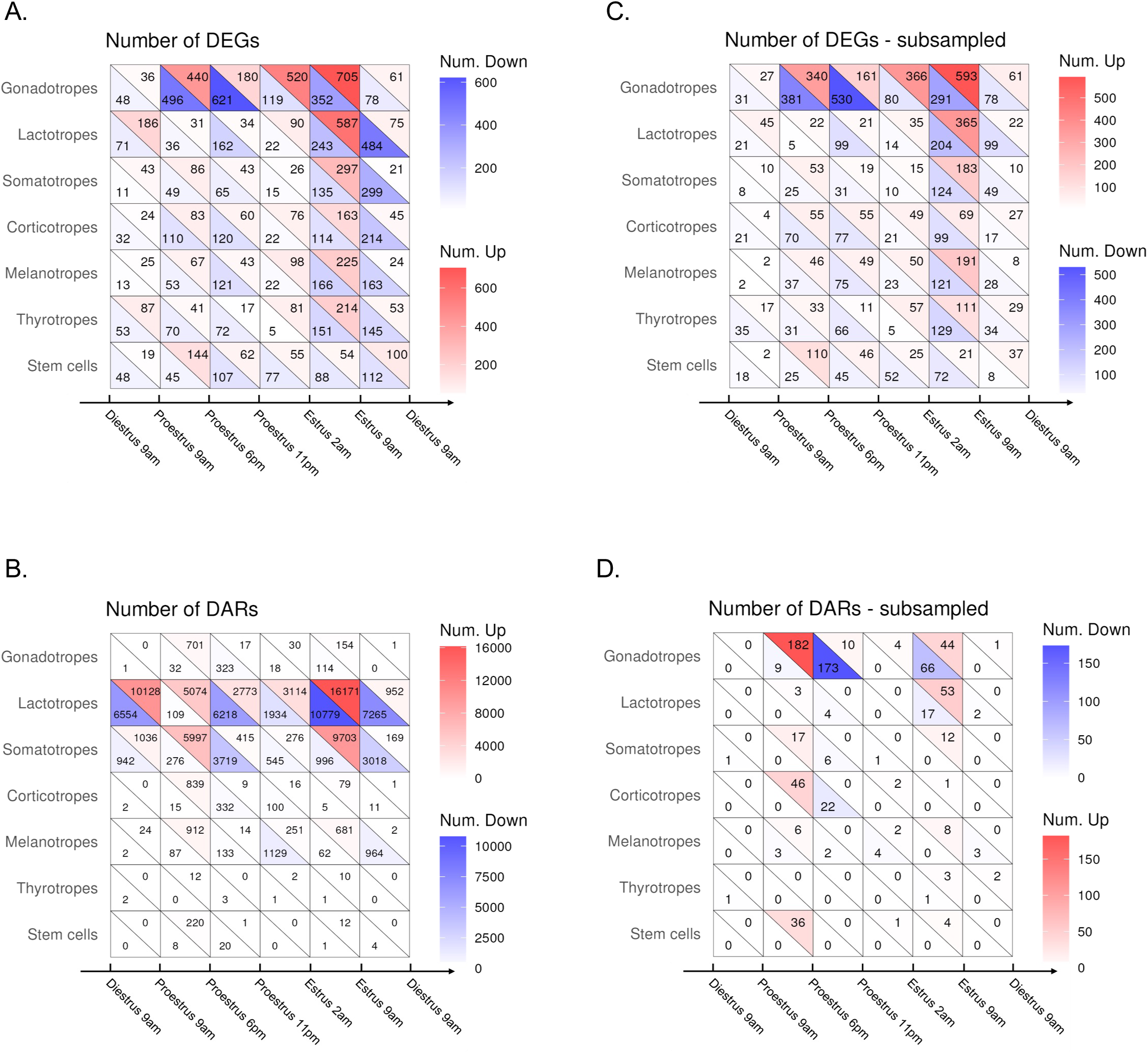
Differential analysis of gene expression and chromatin accessibility. **A,B.** Number of DEGs **(A)** and DARs **(B)** per cell type between two consecutive estrous cycle stages/time points. **C,D**. Number of DEGs **(C)** and DARs **(D)** per cell type between two consecutive estrous cycle stages/time points, with the number of cells being subsampled across cell types and time points, at the exception of diestrus 9 am samples where the number of cells were subsampled across cell types only because only two samples were assayed at that stage. Upregulated genes are in red and downregulated genes are in blue; regions of increased accessibility in red and regions of decreased accessibility in blue.

Because DEG and DAR calling from sc or sn datasets is sensitive to the number of cells studied, we downsampled each cell type to equalize the number of cells in each cell type used for differential analyses. We also equalized the number of cells studied across the different estrous cycle time points except for diestrus 9 am, which was equalized across cell types but had too few cells to equalize across time points (see Methods). While this analysis considerably reduced power to call differential events, especially for DARs, it provided a less biased sense of how many relative events are observed in different cell types at different time points. This equalized analysis showed that while the largest number of DEGs and DARs were detected in the gonadotropes, a large number of regulatory events was also detected in other cell types, particularly in the lactotropes (**Fig. 2C, D**). Another consideration in comparing the estrous cycle programs between cell types was that samples were collected at different times of day, and circadian effects might vary for the different cell types. To control for this, the subsampling analyses were repeated to compare differential expression and differential chromatin accessibility only across the three estrous stages studied at 9 am, namely proestrus 9 am, estrus 9 am, and diestrus 9 am. The results showed a large impact of the estrous cycle stage on all cell types. The number of DEGs was comparable in the gonadotropes and lactotropes, and the largest number of DARs was found in lactotropes (**Fig. S3**). Collectively, these findings support the formulation that all major pituitary cell types go through significant regulatory changes related to the estrous cycle, with the most plasticity seen in gonadotropes and lactotropes (see Discussion). In later sections, we investigated temporal gene clusters, biological processes, integrated GRCs, and the modulation of their activity in the gonadotropes and lactotropes.

### Characterization of cell state dynamics in the gonadotropes during the estrous cycle

Given the structural and functional plasticity displayed by gonadotropes across the estrous cycle (for review, see ^6^), we studied the evolution of cellular state throughout the cycle. The UMAP plot of gonadotropes from all samples showed that samples obtained at the same estrous cycle time point cluster together. Estrus 9 am samples divided into two distinct clusters with all samples from that time point represented in both clusters (**Fig. 3A,B**).

**Figure 3.**
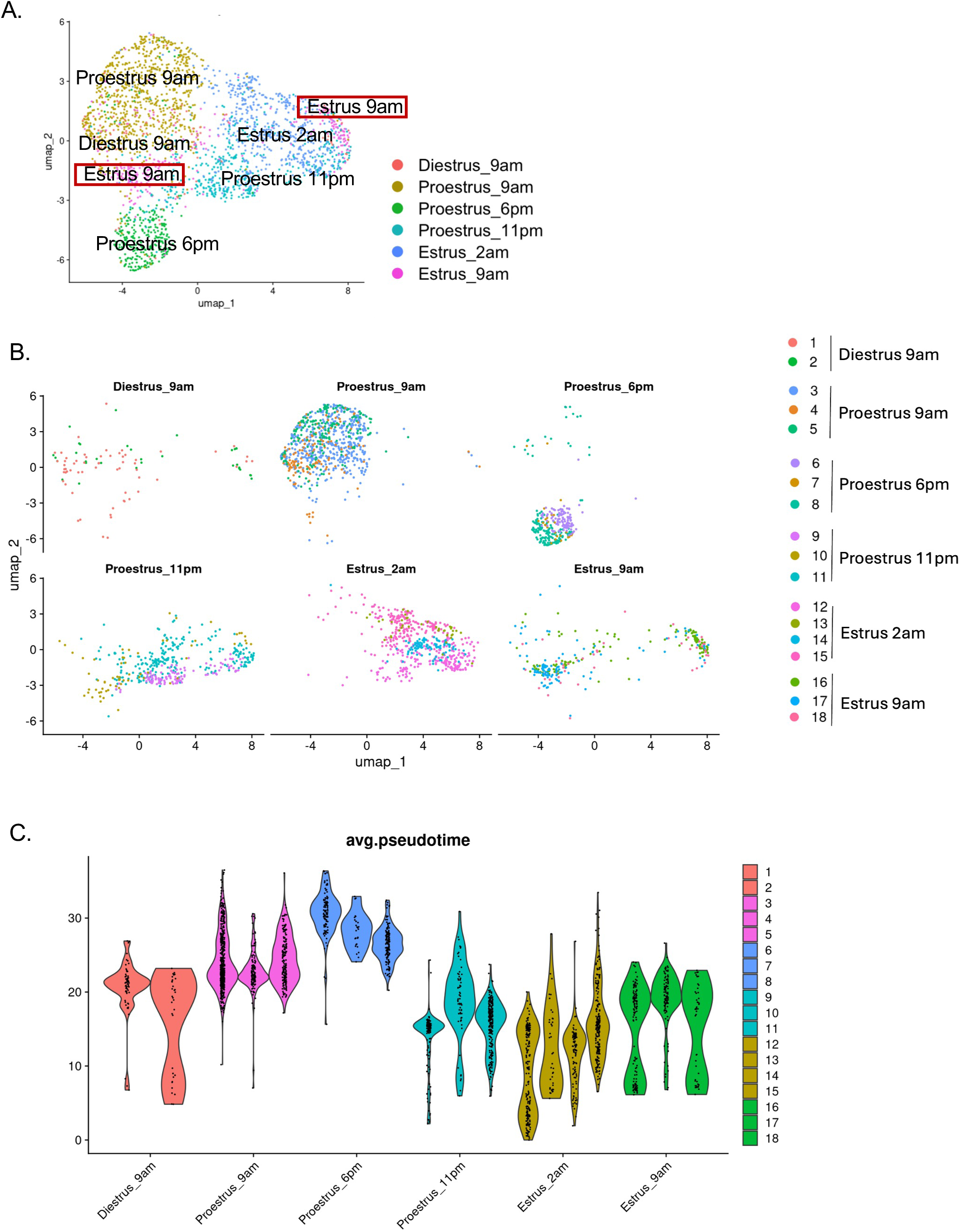
Pseudotemporal trajectory analysis of pituitary gonadotropes during the estrous cycle. **A**. UMAP representation of the gonadotrope cell cluster based on the snRNA-seq data, with labeling by estrous cycle stage/time assignment. Estrous cycle stages are color-coded as indicated. **B**. snRNA-seq UMAP representations of the gonadotrope cell cluster at each of the 6 estrous cycle stages/time points, with labeling by individual animal. Individual animals are color-coded as indicated. **C.** Plot showing the ordering of the gonadotropes by pseudotime. Slingshot trajectory analysis of the gonadotrope cell cluster was inferred from the snRNAseq data, with cells color-coded by animal.

To capture the cell state dynamics of the gonadotropes across the sequential estrous cycle time points, we built a pseudotime trajectory from the snRNAseq datasets using the Slingshot method ^23^. Pseudotime analysis orders individual cells along a virtual timeline based on similarities in their gene expression profiles. This analysis revealed a gradual change in time assignment throughout the cycle, as shown by violin plots (**Fig. 3C**). While most of the samples displayed a predominant pseudotime assignment, several samples (samples 2,12,16,18) showed gonadotropes aggregating at two separate pseudotime positions (**Fig. 3C**). The expression across pseudotime of the 44 genes that are most important for pseudotime assignment is shown in **Fig. S4**. Comparison of expression of the two clusters seen at estrus 9 am identified only *Nnat* as significantly different between the clusters (**Fig. S5**). Eliminating *Nnat* from the analysis, however, did not affect clustering or pseudotime patterns, indicating that the clusters overall showed divergent transcriptome patterns. The lack of characteristic discriminating genes in the two estrus 9 am clusters and the absence of subclusters throughout the estrous cycle suggested that the differences observed represented cell state effects rather than developmentally determined gonadotrope cell subtypes. Thus, while recent lineage tracing studies report two different lineages leading to different gonadotrope subpopulations ^24^, our analysis does not detect distinct gonadotrope subtypes. The pseudotime patterns show some variation between samples obtained from the same estrous cycle stage and time assignment, with some overlap of cells with prior or subsequent cycle stages. This pattern is consistent with the hypothesis that cells rapidly transition from one estrous time point state to another, with some samples showing cells predominantly in one state and others capturing cells before and after transition (see Discussion).

### Temporal dynamics of RNA expression in gonadotropes and lactotropes during the estrous cycle

Alterations in RNA expression and chromatin accessibility in the major pituitary cell types throughout the estrous cycle indicated widespread regulatory changes. To better understand the processes involved, we focused on the gonadotropes and lactotropes, as these cell types showed the largest regulatory effects when cell numbers were normalized (see **Fig. 2**). We first clustered the DEGs from each cell population based on similarity in their temporal expression profiles and then identified the biological processes that were overrepresented among the gene clusters.

In the lactotropes, K-means clustering of the DEGs identified 6 gene clusters with different temporal patterns, which comprised between 74 and 381 genes (**Table S7**). The clusters were labeled according to their patterns of expression change (**Fig. 4A**). Four of the clusters, namely Low estrus 9am-Low diestrus 9am, High proestrus 6pm, Low proestrus 11pm-High estrus 9am, and Low proestrus 11pm-Low estrus 2am, showed large differences between one or more 9 am sample collections, indicating that these estrous cycle changes cannot be entirely accounted for by circadian effects. Pathway enrichment analysis showed that 3 clusters (High proestrus 6pm, Low proestrus 11pm-High estrus 9am, and Low proestrus 9am-High estrus 9am) were dominated by neuronal and synaptic processes, while ribosomal components, endoplasmic reticulum compartment, and mRNA processing were overrepresented in the remaining 3 clusters (Low estrus 9am-Low diestrus 9am, High proestrus 11pm-High estrus 2am, and Low proestrus 11pm-Low estrus 2am; **Fig. S6**).

**Figure 4.**
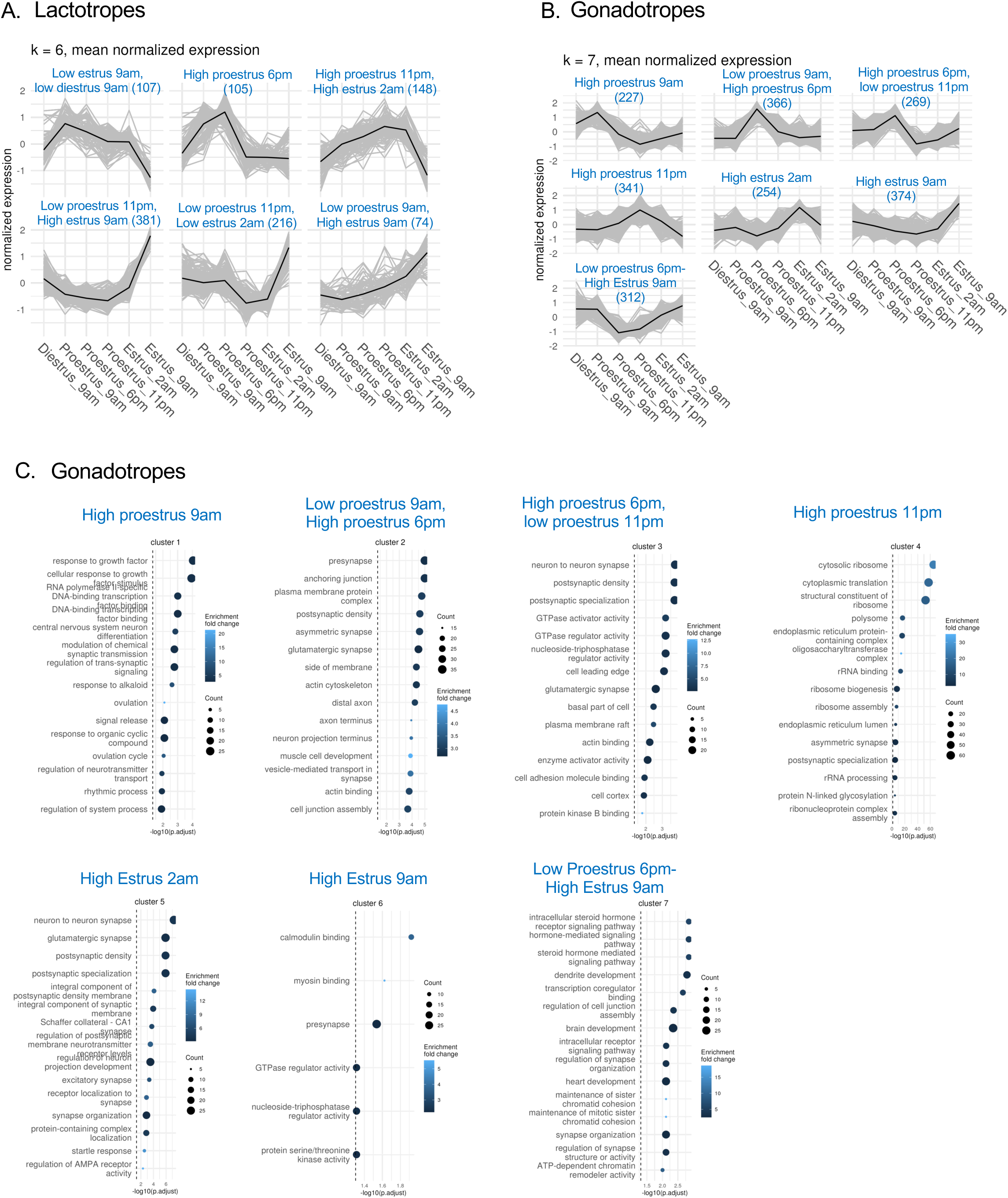
Temporal patterns of gene expression in the lactotropes and gonadotropes. **A.** Identification of 6 clusters of genes in the lactotropes showing distinct temporal patterns across estrous cycle time points. **B**. Identification of 7 clusters of genes in the gonadotropes showing distinct temporal patterns across estrous cycle time points. **C**. Shown are the biological pathways that are over-represented in each of the 7 gene clusters identified in the gonadotropes in **(B).**

In the gonadotropes, K-means clustering identified 7 cluster groups that ranged from 227 to 374 genes (**Table S8**) and showed distinct temporal patterns (**Fig. 4B**). Three of the 7 clusters (Low proestrus 9am-High proestrus 6pm, High proestrus 6pm-Low proestrus 11pm, and High estrus 2am) were enriched for neuronal and synaptic processes. The clusters named High estrus 9am and Low proestrus 6pm-High estrus 9am, which were primarily linked to calmodulin binding and steroid hormone signaling pathway, respectively, were also both associated with synaptic processes. The remaining 2 clusters were annotated to cellular response to growth and ribosomal components (**Fig. 4C**).

These observations support the notion that lactotropes and gonadotropes each transition through different functional states during the estrous cycle, a finding that is in agreement with a recent preprint reporting the sn multiomic analysis of female rat pituitaries across the estrous cycle ^25^.

### Gene regulatory circuitry in gonadotropes and lactotropes during the estrous cycle

Gene regulation is controlled by GRCs, which comprise a triad of a regulatory transcription factor (TF), its *cis*-binding site, and the regulated target gene. GRCs can be summarized at the gene level (multiple TFs and GRCs regulating the same gene) or at the TF level (all GRCs across all genes modulated by a single TF, **Fig. 5A**). To uncover GRCs within the gonadotrope and lactotrope cell types, we used a modification of the Multiome Accessibility Gene Integration Calling and Looping (MAGICAL) framework ^16^ for analysis of the sn multiome data. MAGICAL identifies a GRC activity that varies across cells and conditions in each cell type through mapping a regulatory domain with a chromatin accessibility change to altered mRNA expression of a target gene and of the regulatory TF (**Fig. 5B**). In the original implementation of MAGICAL, cells were analyzed by cell type using the average of RNAseq and ATACseq data for each cell type. To increase statistical power and leverage the matched same-cell multiome data that we generated, rather than apply MAGICAL to the average values of the lactotropes or gonadotropes in each sample as originally implemented, we grouped the lactotropes or gonadotropes into metacells using the SEACells method on the snRNAseq data (^26^; see Methods and **Fig. S7**).

**Figure 5.**
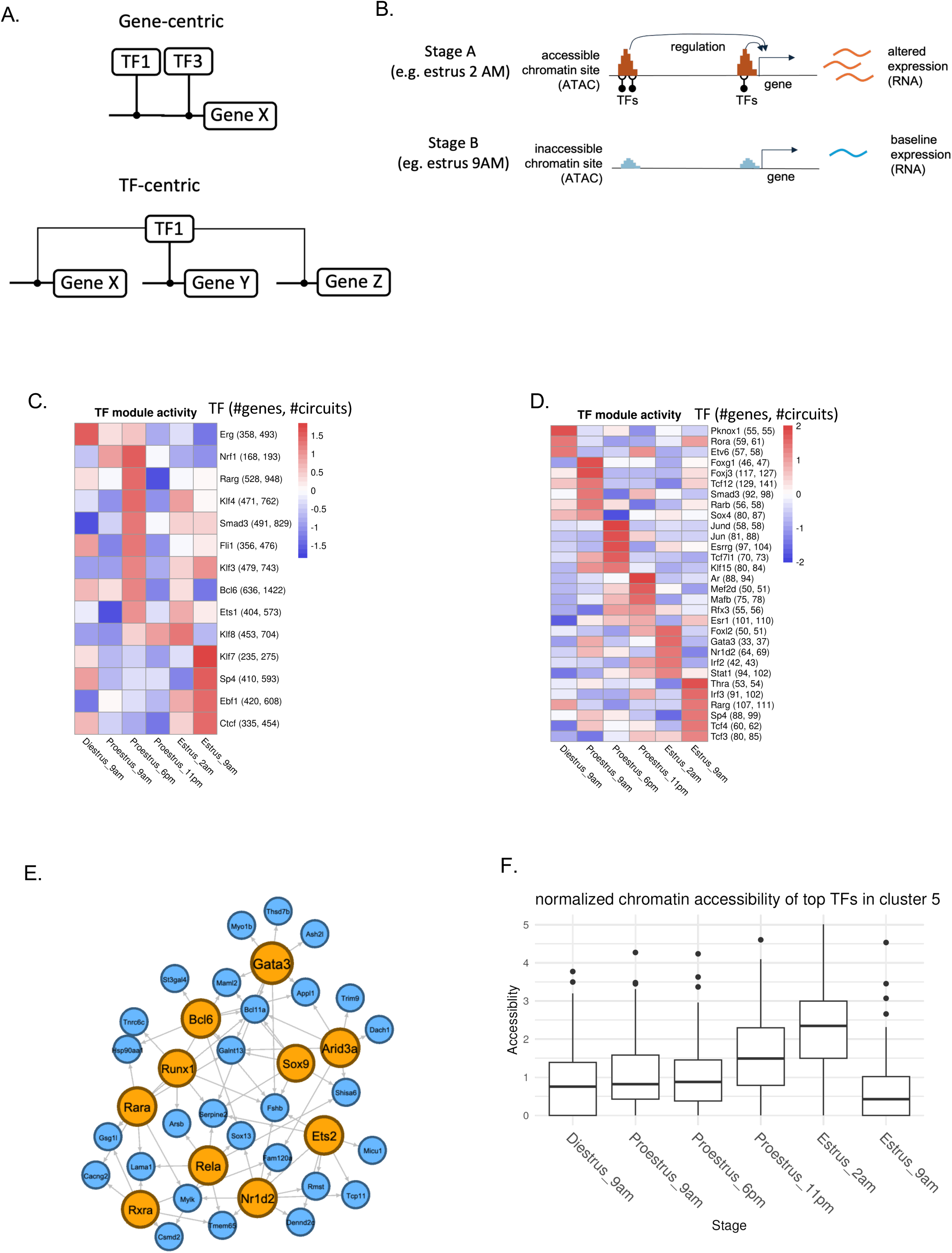
Identification of gonadotrope and lactotrope GRCs across the estrous cycle. **A**, **B**. Schematic representation of a GRC. The GRC triad consists of a transcription factor (TF), its *cis*-binding site, and the regulated target gene. **A**. *Top*, Gene-centric circuit, where multiple TFs regulate a given gene. **A**. *Bottom*, TF-centric circuit, where multiple genes are regulated by a specific TF. **B**. Example of a GRC where the TF differentially regulates a specific target gene between two consecutive estrous cycle time points; the GRC exhibits changes in: i) RNA expression of the target gene, ii) chromatin accessibility of the TF-binding region on the target gene, iii) RNA expression of the TF-encoding gene. **C,D**. Heatmaps of the average activity per estrous cycle time point for the top TF modules identified in the lactotropes (**C**) and the gonadotropes (**D**); for details, see Methods. Each TF module is comprised of a regulatory TF, its target genes, and the total number of GRC triads/circuits it is involved in, as indicated on each row of the heatmap. **E**. Network analysis of the top 10 regulatory TFs in the Peak estrus 2am cluster identified in the gonadotropes. Orange-colored circles represent the top regulatory TFs between estrus 2am and estrus 9am; blue-colored circles signify the genes targeted by these TFs and showing differential expression between estrus 2am and estrus 9am. **F**. Average chromatin accessibility levels measured in the TF-binding regions of target genes for the top 10 regulatory TFs identified in the Peak estrus 2am cluster.

We initially identified the lactotrope TF modules showing the highest levels of activity when comparing all samples and estrous cycle time points (aggregate TF module activity; see Methods for details). **Fig. 5C** illustrates the activity of those top TF modules at each estrous cycle time point (see **Table S9** for a complete list of all GRCs). The TFs regulating the most genes included Nrf1 at proestrus 9am; Rarg, Klf4, and Smad3 at proestrus 6pm; Klf8 at proestrus 11pm and at estrus 2am; Klf7 at estrus 9am. We noticed some overlap in TF module activity between time points; for instance, Nrf1 module activity was high at both proestrus 9am and proestrus 6pm, and Klf3 module activity was high at both proestrus 6pm and estrus 9am. Hence, while regulatory circuits may show changes in activity at specific time points, some of them can exhibit similar activity at different time points.

Using the same approach for the gonadotropes, we next identified the gonadotrope GRCs showing the highest levels of activity when comparing all samples and estrous cycle time points. **Fig. 5D** depicts the activity for the top TF modules at each estrous cycle time point. A complete list of all GRCs is provided in **Table S10**. Top TFs included: Foxg1, Foxj3, Smad3 at Proestrus 9am; Jund, Jun, Esrrg at proestrus 6pm; Ar and Mef2d at proestrus 11pm; Foxl2 and Gata3 at estrus 2am; Thra and Irf3 at estrus 9am. Interestingly, among the top lactotrope and gonadotrope TF modules, the regulatory TFs Sp4, Smad3, and Rarg were shared between the two cell types (**Fig. 5C,D**). While Sp4 module activity was the highest at Estrus 9am in both lactotropes and gonadotropes, the activities of the Smad3 and Rarg modules peaked at different time points in each cell type. Rarg was previously found to be expressed in Rathke’s pouch and the infundibulum (pituitary stalk) during early pituitary development ^27^. In the gonadotropes, Smad3 is known to mediate activin-induced *Fshb* gene expression ^28^, while TGFβ/Smad3 signaling contributes to the inhibitory effects of dopamine on lactotrope cell proliferation and PRL secretion ^29,30^. When examining the target gene overlap between the lactotrope and the gonadotrope Smad3 modules, we identified 12 common genes, which showed enrichment for synaptic processes (**Fig. S8**). The lactotrope-specific target genes of the Smad3 module were also enriched for synaptic processes, whereas processes related to reproductive development were overrepresented among the gonadotrope-specific target genes. Overall, these results support the formulation that circuit activity - with its associated TF expression level and altered chromatin accessibility at the TF binding site - is a major driver of gene expression variation across the estrous cycle.

To explore gene regulatory mechanisms contributing to the pre-ovulatory surge in gonadotropins ^4^, we focused on the 254 DEGs comprising the peak estrus 2am cluster identified in the gonadotropes (**Fig. 4B**). The ten TFs (Gata3, Bcl6, Sox9, Arid3a, Runx1, Rara, Ets2, Rela, Rxra, and Nr1d2) that regulated the largest number of genes at estrus 2am are shown along with their target genes in the network schematic in **Fig. 5E**. Many of the target genes were regulated by multiple TFs. As expected, the overall chromatin accessibility at the *cis*-binding sites of those target genes was greatest at peak estrus 2 am (**Fig. 5F)**. Notably, the *Fshb* gene was among the target genes whose expression was significantly altered at estrus 2am, and its regulatory TFs included Runx1, Bcl6, Gata3, Sox9, Nr1d2, and Ets2 (**Fig. 5E** and **Fig. S9**).

### Role of ETS2 in modulating Fshb gene expression

We sought to investigate further the regulatory control of the *Fshb* gene. We recently reported the characterization of 4 upstream enhancers in the murine *Fshb* gene, referred to as Enh1 through Enh4. Enh3 and Enh4 were found to be most important for *Fshb* gene regulation by activin *in vitro* ^31^. As described in the GRC analysis in the previous section, the transcription factor ETS2 is one of the major drivers of gene regulation at estrus 2am and is implicated in *Fshb* regulation (See **Fig. 5E**). Enh4, which contains the binding site for the ETS2-*Fshb* GRC, shows the greatest changes in chromatin accessibility during the cycle (**Fig. 6A** and **Fig. S9**). *Fshb* mRNA levels rise dramatically beginning at proestrus 11 pm, peaking at estrus 2 am, and then returning to baseline at 7 hours later at estrus 9 am (**Fig. 6B**). The dramatic increase in *Fshb* transcript levels observed on proestrus 11 pm and then on estrus 2 am is consistent with earlier studies in rats ^32^. *Ets2* mRNA levels and ETS2 inferred activity also peak at estrus 2 am (**Fig. 6B** and **Fig. S9**). However, the TF mRNA and inferred activity are still at baseline at proestrus 11 pm, an estrous cycle timepoint at which *Fshb* is already rapidly increasing. The maximal chromatin accessibility for the Enh4 ETS2 binding site is seen at estrus 2 am (**Fig. 6C**). These *Fshb* and *Ets2* RNA expression and chromatin accessibility dynamics, with *Ets2* increases lagging those of *Fshb,* suggest that ETS2 functions as a repressor of *Fshb* at the Enh4 site and contributes to its rapid return to basal levels by estrus 9 am.

**Figure 6.**
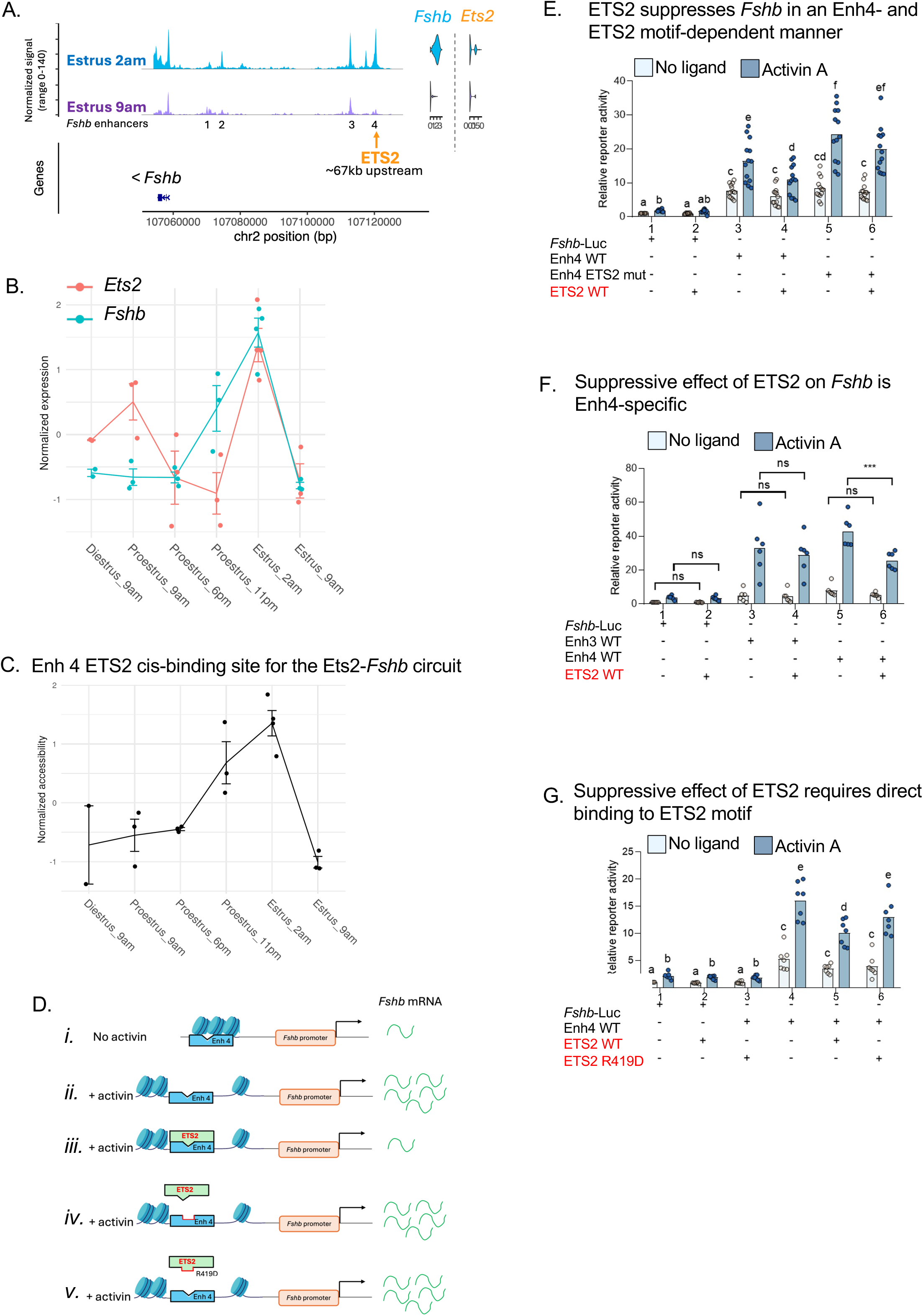
Experimental validation of the predicted ETS2-*Fshb* circuit. **A.** Shown are the chromatin accessibility tracks of the *Fshb* gene and 5’ flanking region at estrus 2 am and at estrus 9 am (*Left*) along with *Fshb* and *Ets2* RNA expression levels at the same stages (*Right*). The position of the ETS2 binding site within Enh4 is signified by an orange arrow. Chromatin accessibility and RNA expression signal were normalized. Presented at the bottom of the figure is a schematic view of the genomic region holding the *Fshb* gene and its 5’ flanking region, with genomic coordinates. **B,C.** Line graphs illustrating the coordinated RNA expression of *Ets2 (in red)* and *Fshb (in blue;* (**B**)), and chromatin accessibility at the ETS2 binding site (**C**) across the estrous cycle. **D**, Schematic summary of the key findings obtained in LβT2 cells in previous work and in panels **E-G**. *(i)* In the absence of activin, the Enh4 domain is less accessible ^31^ and *Fshb* mRNA expression is low ^33,34^. *(ii)* In the presence of activin, Enh4 is highly accessible (Jin 2025) and *Fshb* mRNA expression is high ^33,34^. *(iii)* ETS2 exerts an inhibitory effect on activin-induced *Fshb* mRNA expression, likely through binding to the ETS2 binding motif in Enh4 (see panels **E-G**). *(iv)* The inhibitory effect of ETS2 is abolished when the ETS2 binding motif is disrupted/mutated (see panel **E**). *(v)* The inhibitory effect of ETS2 is abolished by mutation in the DBD of ETS2 that blocks DNA binding(see panel **G**). (**E-G**) LβT2b cells were transfected with 225 ng/well of one of the indicated reporter constructs, with or without 5 ng/well of one of the expression constructs (pcDNA3.0 was used to equalize total DNA), indicated in red. Data were log-transformed and analyzed by two-way ANOVA followed by Holm-Šidák test. (**E,G**) Transfected cells were treated with either no ligand (vehicle) or 0.05 nM Activin A for 6 hours. N = 13 (**E**) and N = 7 (**G**) independent experiments, as specified by the number of individual points. Different letters indicate statistical significance. **F.** Transfected cells were treated with either no ligand or 0.5 nM Activin A for 6 hours. N = 6 independent experiments. ***, P<0.001; ns, not significant.

We next performed experiments to investigate the predicted regulatory effects of the Enh4 ETS2-*Fshb* GRC using reporter assays in the LβT2 mouse gonadotrope-like cell line. As illustrated schematically in **Fig. 6D**, our previous bulk ATACseq analyses in LβT2 cells showed that, in the absence of activin A, the Enh4 region/ETS2 motif was compacted, whereas in the presence of activin A it became highly accessible ^31^. In addition, earlier LβT2 cell studies found that *Fshb* mRNA expression is strongly induced by activin A ^33,34^. Accordingly, in our ETS2-*Fshb* validation experiments, we compared the effects of ETS2 overexpression on the activin A-induced transcriptional activity of *Fshb* promoter-reporter constructs with and without enhancers.

We first compared the effects of ETS2 overexpression on a minimal *Fshb* promoter (-1990/+1) construct referred to as *Fshb*-luc (and previously described in ^35^) relative to an *Fshb* promoter-reporter construct including Enh4 region, Enh4 WT (**Fig. S11A** and ^31^). To overexpress ETS2, we co-transfected a FLAG-tag expression vector encoding the wild-type ETS2 TF (ETS2 WT; **Fig. S11A**). As shown in **Fig. 6E**, *Fshb* promoter activity alone was very modestly induced by activin A (lane #1), and overexpression of ETS2 WT had no detectable effect on this induction (#2 vs. #1). Insertion of Enh4 WT led to a significant increase in activin-induced promoter activity compared to *Fshb*-luc (#3 vs. #1), a result that was consistent with the enhancer activity of Enh4 ^31^. Co-transfection with both Enh4 WT and ETS2 WT resulted in a significant reduction in activin-induced transcriptional activity relative to Enh4 WT alone (#4 vs. #3), consistent with ETS2 acting as a repressor at Enh4. To determine whether the effect of ETS2 depended on the ETS2 binding site present in Enh4, we introduced two point mutations in the ETS2 binding motif that were predicted to cause a 5-fold decrease in ETS2 binding probability (see Methods). The binding site mutation increased the level of *Fshb* activation, consistent with the mutations interfering with the effects of endogenous ETS2. Suppression of the activin-induced reporter activity by ETS2 overexpression relative to this baseline was not seen with this Enh4 Ets2 mutation construct (**Fig. 6E**). These results are consistent with the hypothesis that ETS2 binds at Enh4 to attenuate activin-stimulated *Fshb* expression.

We next evaluated the enhancer specificity of the observed ETS2 repressor activity. We compared the effects of ETS2 overexpression on activin-induced *Fshb* promoter activity in the presence of either Enh4 (Enh4 WT) or Enh3 (Enh3 WT). Because the Enh3 WT construct required a higher concentration of activin A to elicit a significant increase in activin-induced transcriptional activity, we used a 10-fold higher activin concentration in these experiments for both constructs. In contrast to the suppression seen with the Enh4 WT construct, ETS2 overexpression had no effect on activin-stimulated induction of the Enh3 WT construct (**Fig. 6F)**. These observations further support the hypothesis that ETS2 exerts an inhibitory effect on activin-induced *Fshb* promoter activity via Enh4. To provide further evidence that the ETS2 suppression depended on ETS2 binding to Enh4, we compared the effect of overexpression of ETS2 WT on activin-induced promoter activity in Enh4 WT to that of overexpression of ETS2 R419D, in which one of the two key DNA-contacting arginine residues (R419) was mutated to impair DNA binding activity ^36–38^ (see Methods). The mutant ETS2 did not significantly suppress activin induction of the Enh4 WT *Fshb* construct (**Fig. 6G**, see **Fig. 6D)**. We also evaluated the effects of wild-type and mutant ETS2 on endogenous *Fshb* expression. The LβT2 cells were transfected with either with an HA-tag expression vector encoding ALK5TD, a constitutively active form of the type I receptor, ALK5 ^39^, or with a combination of ALK5TD and a FLAG-tag expression vector expressing ETS2 WT. Transfection with ALK5TD-HA alone caused a substantial increase in *Fshb* transcript levels relative to transfection with empty expression vector (pcDNA3.0; **Fig. S11B**), reflecting the stimulatory effect of activin-like signaling on endogenous *Fshb* gene expression. Co-expression of ETS2 WT significantly decreased ALK5TD-stimulated *Fshb* mRNA expression, thus demonstrating an inhibitory effect of ETS2 on ‘activin’-induced expression of the endogenous *Fshb* gene. Conversely, when ETS2 R419D was co-expressed with ALK5TD, the *Fshb* stimulation by activin signaling was not affected (**Fig. S11B**). These results further support the inhibitory effect of ETS2 on activin-induced *Fshb* gene expression.

Collectively, these studies provide experimental support for the computationally inferred regulatory role of ETS2 in *Fshb* gene expression during the murine estrous cycle. In light of the *in vitro* suppressive effect of ETS2 on activin-induced *Fshb* gene transcription, we propose that the rise in *Ets2* mRNA at estrus 2 am leads to an increase in ETS2 expression that is likely to contribute to the subsequent decline in *Fshb* mRNA at estrus 9 am.

## Discussion

Our analysis of carefully staged mice characterized the changes in gene expression and chromatin accessibility occurring throughout the estrous cycle within each individual nucleus across pituitary cell types. The study revealed that a high level of transcriptomic and epigenetic plasticity occurs during the estrous cycle, not only in gonadotropes but also across other pituitary cell types. The use of sn multiome data allowed for the computational identification of the GRC triads (gene, regulatory site, transcription factor) underlying these gene expression changes. Experimental confirmation of the predicted role of ETS2 acting at a specific upstream enhancer to regulate *Fshb* expression in gonadotropes supports the value of this multiomic dataset and its analysis for gaining mechanistic, cell type-level insight into the regulatory processes underlying pituitary plasticity during the estrous cycle.

Pituitary cell populations have been postulated to undergo restructuring in response to physiological signals using various cellular and molecular mechanisms (for review, ^40,41^). At the cellular level, these mechanisms can include alterations in the proportion of responding cells, in the amount of hormone secreted per cell, and in the number of cells from the same population, presumably through cellular proliferation, transdifferentiation (i.e., the transformation of cells from one hormone-producing cell type to another), and differentiation of stem/progenitor cells. At the molecular level, epigenetic and transcriptomic remodeling and their impact on protein expression and protein activity are implicated. We found widespread gene expression and chromatin accessibility changes at each time point through the estrous cycle. However, our analysis did not identify changes in defined subpopulations of gonadotropes or changes in cell proportions indicative of proliferation or transdifferentiation. These results are consistent with a high level of cell plasticity in the gonadotrope and other cell types during the cycle, with the changes observed reflecting dramatic alterations in cell state.

The pseudotime trajectory analysis indicates that at a given cycle stage and time point, gonadotrope cells can co-exist in different states (see **Fig. 3C**). This co-existence of different cell states may reflect a rapid transition of individual cells occurring so that some samples are capturing cells on both sides of this transition. In earlier sc studies of the LβT2 gonadotrope-like cell line, we found that gene expression changes in response to increasing GnRH stimulation were bimodal, with higher concentrations of GnRH leading to an increased probability that any given cell would convert to a high gene activity state ^42^. Based on the present trajectory results, we speculate that the overall responses during the estrous cycle are similar, with individual cells showing a rapid transition between distinct cell gene expression states.

The large number of genes showing regulation during the cycle is consonant with the gonadotrope’s prominent role in estrous cycle control. Additionally, we observe a large number of regulatory events in other cell types. To control for the effects of differences in cell numbers detected when comparing cell types, cell populations were subsampled for differential analysis. This demonstrated that the number of DEGs is greatest in gonadotropes (2,939 total DEG events) with lactotropes having the second largest number (952 total DEG events; see **Fig. 2C**). A significant number of DEGs were also identified in other major cell types. For example, in the estrus 2 am to estrus 9 am comparison, the subsampled gonadotropes had a total of 884 DEG events, while the subsampled lactotropes, melanotropes, somatotropes, thyrotropes, and corticotropes showed 64% (569 DEG events), 35% (312 DEG events), 35% (307 DEG events), 27% (240 DEG events), and 19% (168 DEG events) of this number, respectively. Importantly, by including samples from three distinct estrous stages (P, E, and D), which were all collected at 9 am, we were able to exclude circadian influences on pituitary cell type patterns of differential expression and chromatin accessibility. Lactotropes were previously reported to exhibit changes in their responsiveness to hypothalamic stimuli such as dopamine and oxytocin during the estrous cycle ^43,44^. Previous rat studies also described expression changes in the somatotropes during the estrous cycle ^45–48^. Overall. our analysis demonstrates that all the main pituitary cell types show considerable plasticity during the estrous cycle. The differential RNA expression signal that we noted in lactotropes during the estrous cycle is also compatible with earlier rat studies that reported expression changes in primary lactotrope cells following estrogen exposure or *in vivo* ^49,50^.

In the gonadotropes, pathway analysis of the DEGs showed enrichment for endocrine-related as well as neuronal processes. The enrichment for neuron-related pathways in 5 of the gonadotrope temporal trajectory clusters (and in 3 of the lactotrope clusters) are consistent with the molecular similarities between pituitary and neuronal ontogenesis (^51^; for review, see ^52^). Le Cicle et al. demonstrated that the Neurod1/4-Ntrk3-Src activation pathway regulates the motility of immortalized αT3-1 gonadotrope-like cells and of gonadotropes during mouse pituitary development ^51^, and previous studies showed that NeuroD1 promotes cell motility in human neuroblastoma and bronchial epithelial cell lines ^53,54^. Moreover, earlier work indicated that GnRH induces cell movement in the pituitary ^55^. One possibility is that the regulation of some genes implicated in neuronal processes, e.g. Neurod1/4 and Ntrk3, may contribute to cellular state changes as well as cell migration during the estrous cycle.

GRCs, which consist of a TF, a *cis*-binding site with which it interacts, and a target gene whose transcription is altered, represent the major mechanisms controlling how a chromatin region and TF inputs are converted into mRNA expression level outputs ^56^. Our snRNAseq/ATACseq multiome data, coupled with the MAGICAL framework, enabled us to build a high-resolution map of the transcriptional regulatory circuitry underlying gene expression changes occurring in the gonadotrope and lactotrope cell types throughout the estrous cycle. Several thousands of GRCs were inferred computationally. Among the regulatory TFs driving GRCs in the gonadotropes, ESR1 (or estrogen receptor alpha) is known to mediate the estrogen negative feedback on the gonadotrophs, as gonadotroph-specific knockout of the *Esr1* gene in mice previously resulted in high levels of serum LH and elevated *Lhb* subunit gene expression^57^. Another regulatory TF, Smad3, is involved in the regulation of the *Fshb* and *Gnrhr* genes by activins (for review, see ^58–60^). GATA3 was previously implicated in activating *CGA* gene expression both in the pituitary and in the placenta ^61^. GATA3 is a paralog of GATA2, a TF involved in the determination of the gonadotrope and thyrotrope cell types during mouse pituitary development ^62^.

Our analysis identifies an *Fshb* GRC with ETS2 acting at upstream enhancer 4. The temporal dynamics of *Ets2* and *Fshb* changes between proestrus 6 pm and estrus 9 am are consistent with the idea that the activity of this circuit attenuates *Fshb* gene synthesis and contributes, along with increases in inhibin B, to the rapid decrease in *Fshb* expression between estrus 2 am and 9 am. Our studies in the LβT2 gonadotrope-like cells are consistent with the model, showing that ETS2 attenuates activin-induced *Fshb* gene expression through binding to Enh4. According to previous work, ETS2 interacts with the TGFβ-Smad pathway (^63^; for review, see ^64^). We previously reported that activin stimulates recruitment of SMAD2/3 and FOXL2 to Enh4 ^31^. Overall, these results raise the possibility that the repressive effect of ETS2 may result from it interfering with the activity of activin-stimulated TFs at Enh 4 (for review, see ^58,65^).

In conclusion, our study outlines the dynamic transcriptomic and epigenomic changes occurring in the murine pituitary across the estrous cycle at sn resolution. We characterized cellular state dynamics in the gonadotrope population, identified gonadotrope and lactotrope GRCs, and provided experimental validation for the ETS2-driven GRC as a mechanism in the regulation of *Fshb* gene expression in the gonadotropes. Our analysis provides a framework for the description of cell state and GRC dynamics in the mouse pituitary across the reproductive cycle.

### Limitations of the study

Estrous cycle staging can be difficult and variability between individual animals can be high. We have addressed these issues using multimodal staging and multiple animals for each timepoint. While this is a massive study in comparison with any comparable efforts, the resolution is nonetheless limited by the number of animals at each timepoint. Additionally, our study is limited to 6 estrous cycle timepoints. Analysis of additional timepoints and large sample numbers would provide an even higher resolution map of estrous cycle gene regulatory dynamics. Our analysis is limited to gene expression and chromatin accessibility and does not capture other omic layers such as histone modification state, DNA methylation, and protein expression. Future studies that overlay additional modalities on the map that we have generated will provide further insight into the regulatory control mechanisms underlying the estrous cycle.

## Resource availability

### Lead contact

Further information and requests for resources and reagents should be directed to and will be fulfilled by the lead contact, Frederique Ruf-Zamojski.

### Materials availability

This study did not generate new materials.

### Data and code availability

- The sn multiomic datasets generated in the present study have been deposited and will be available upon request.
- the sn estrous multiomics atlas can be browsed via a web-based portal accessible at https://rstudio-connect.hpc.mssm.edu/snpit_estrous_browser/
- Any additional information required to reanalyze the data reported in this paper is available from the lead contact upon request.

## Acknowledgments

This work was supported by funding from NIH grant NIH DK46943 to SCS, and Canadian Institutes of Health Research (CIHR) project grants PJT-169184, -191766, and -195832 to DJB. We acknowledge the New York Genome Center and New York University for sequencing. This work was supported in part through the computational resources and staff expertise provided by Scientific Computing at the Icahn School of Medicine at Mount Sinai.

## Author Contributions

FRZ, SCS, DJB designed and coordinated the study. LO, AA, XZ, EB performed animal experiments and collected animal samples. FRZ, NM, and GS performed NGS experiments. WSC, ZZ, MZ, EZ, GRS curated the data/performed the QC analysis. WSC, ZZ, OGT, and EZ performed the bioinformatics analysis. YFL prepared DNA constructs, performed cell culture and protein quantification experiments. FRZ, SCS, DJB, and HP interpreted the data. FRZ and HP drafted the original manuscript draft. FRZ, ZZ, WSC, SCS, DJB, and JLT gave comments and edited the manuscript. All authors reviewed the manuscript.

## STAR Methods

### Key resources table

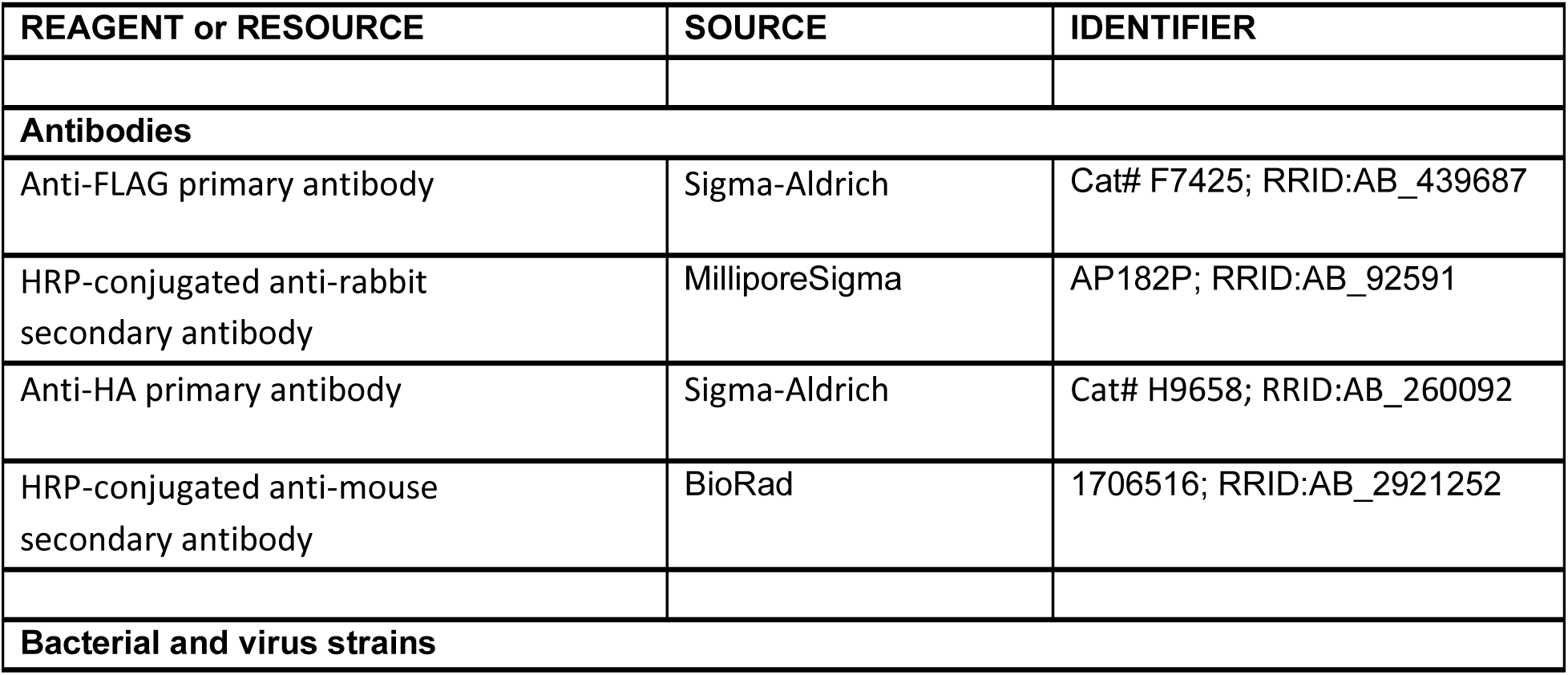
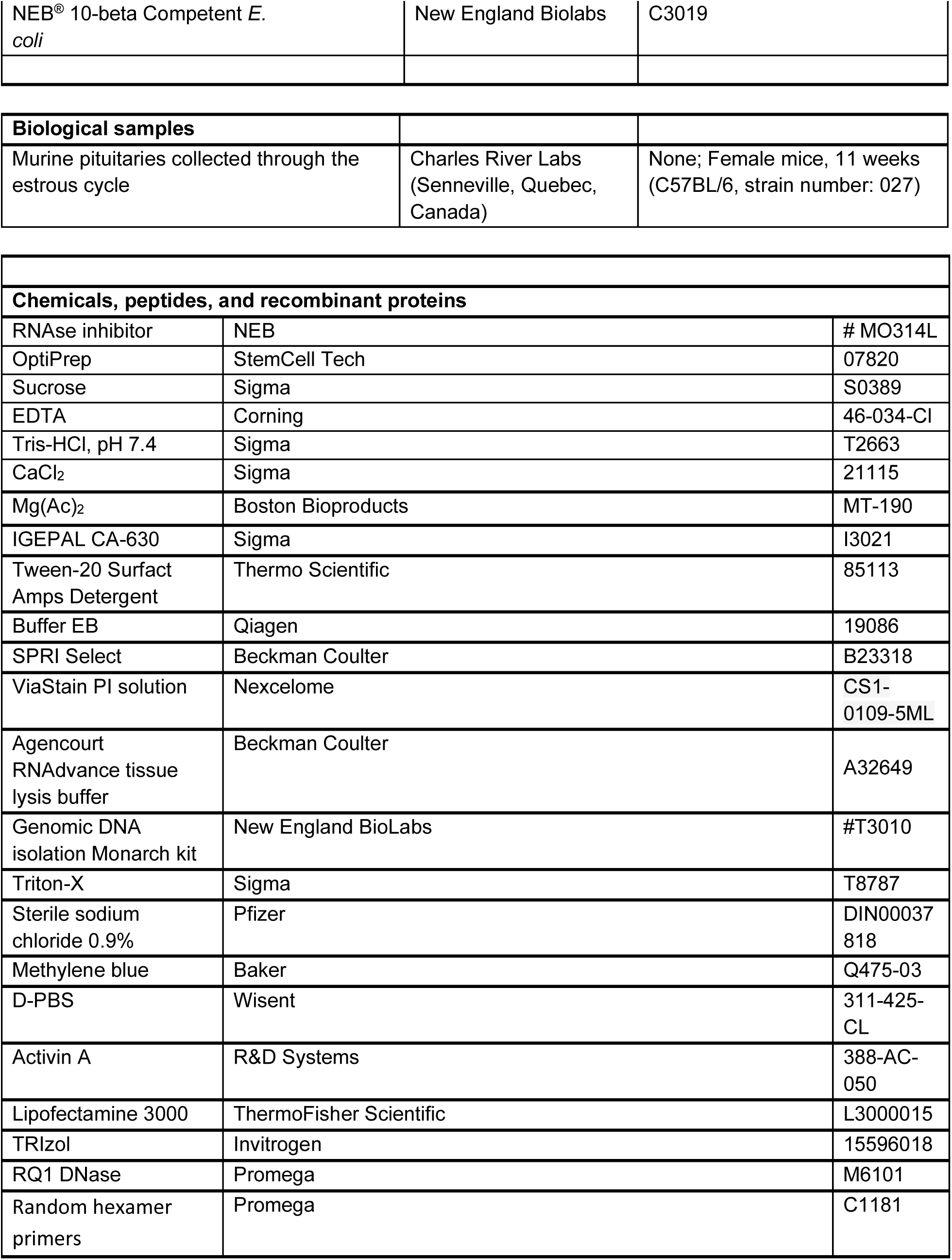
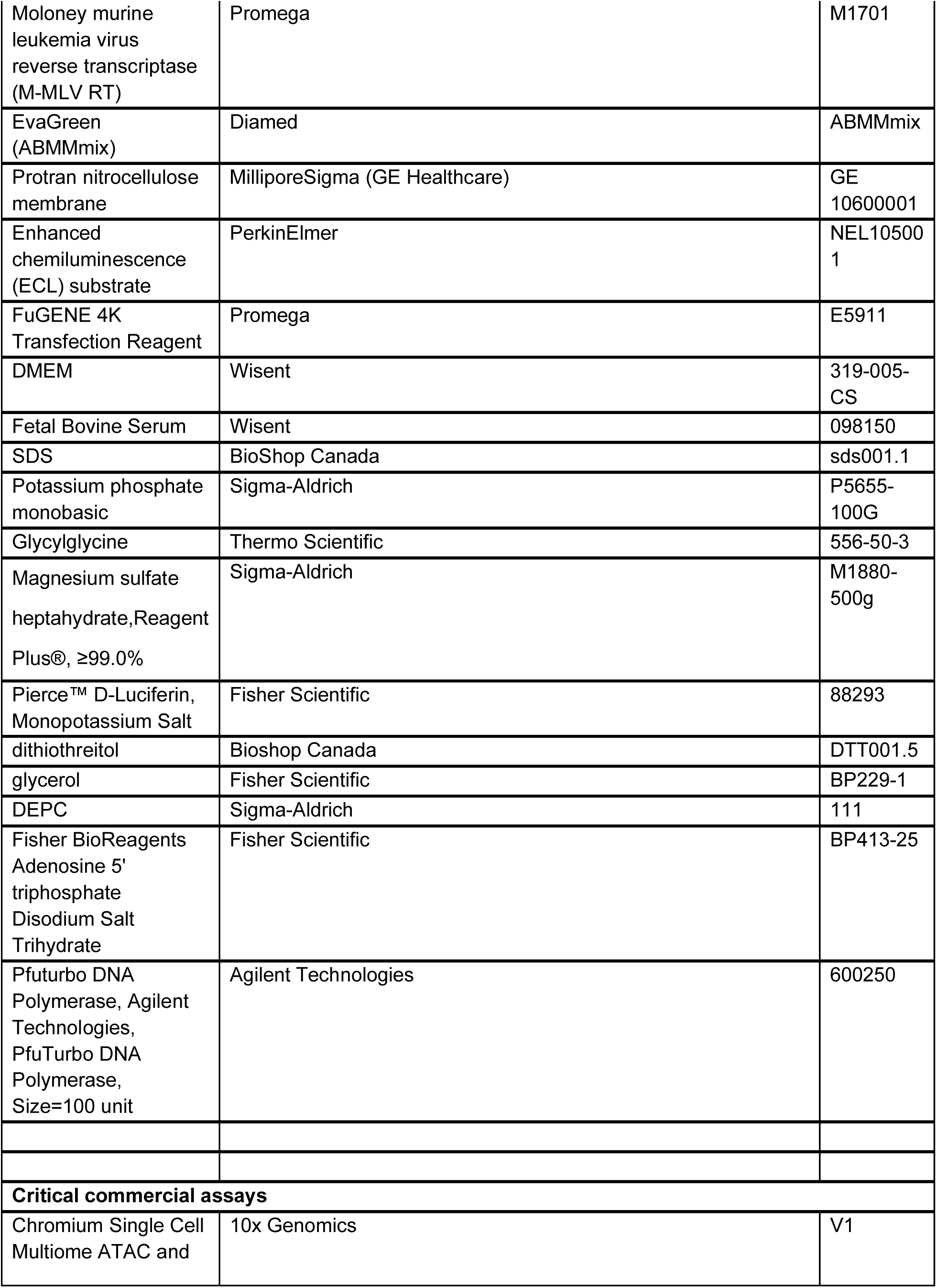
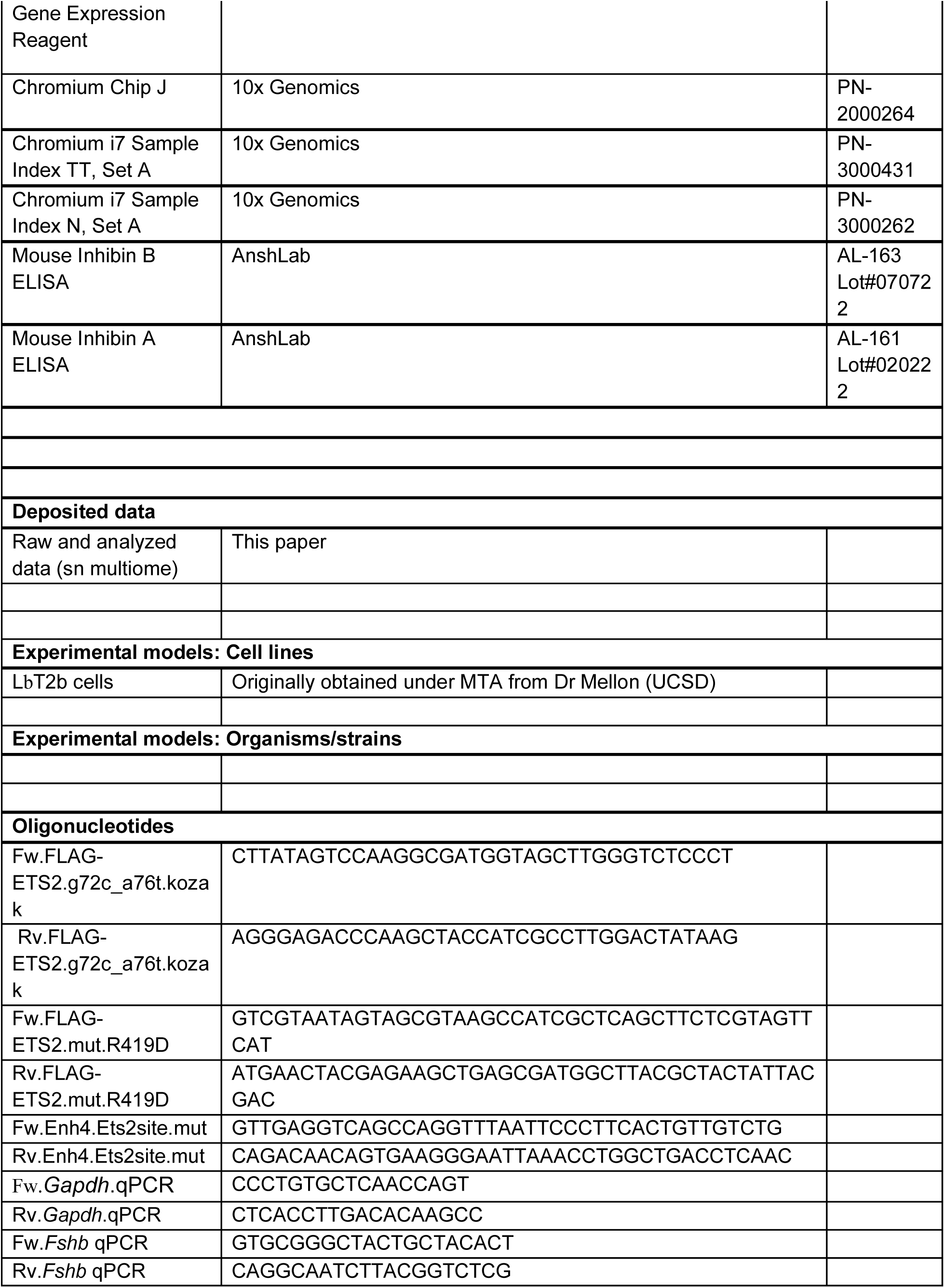
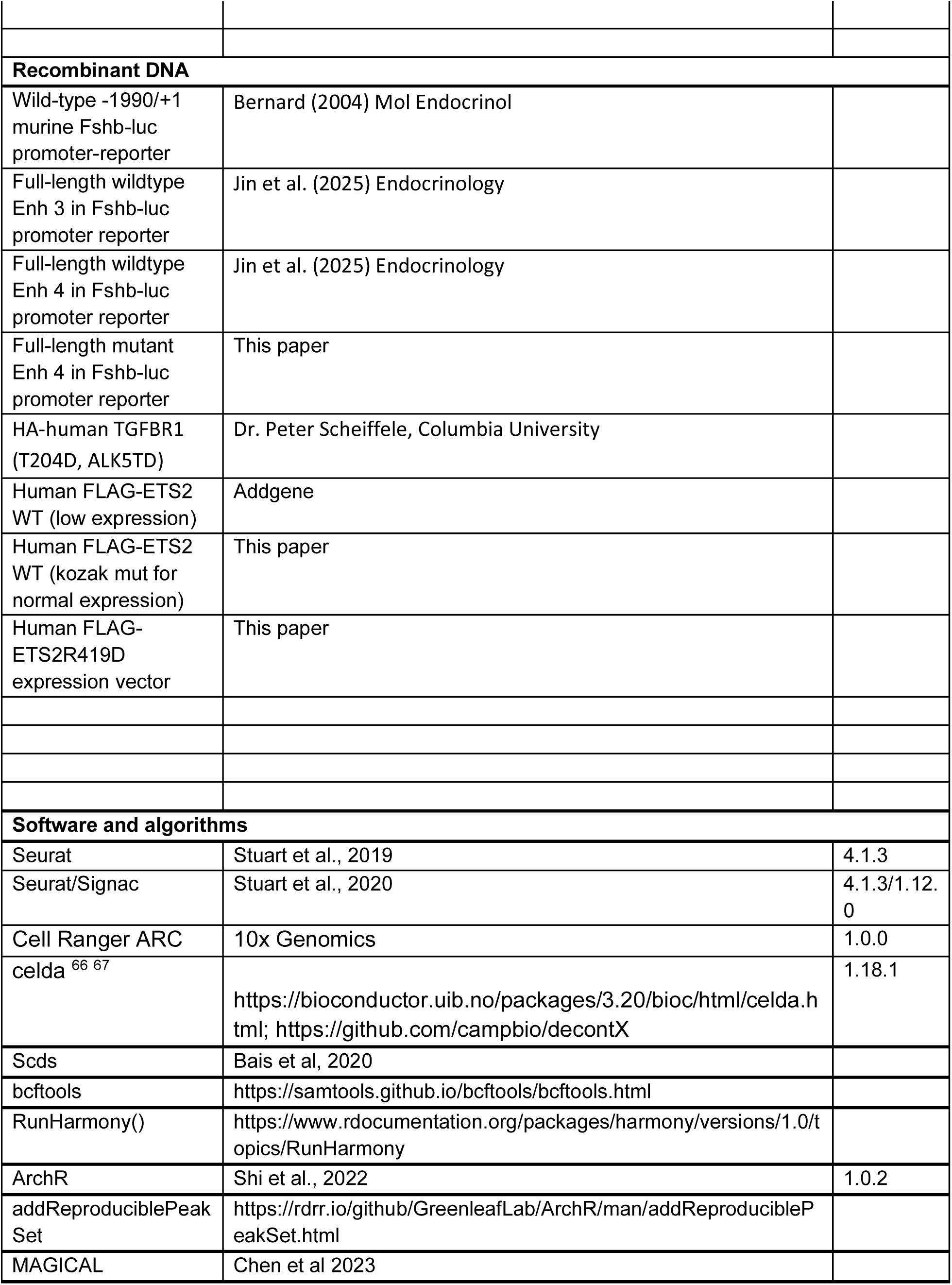
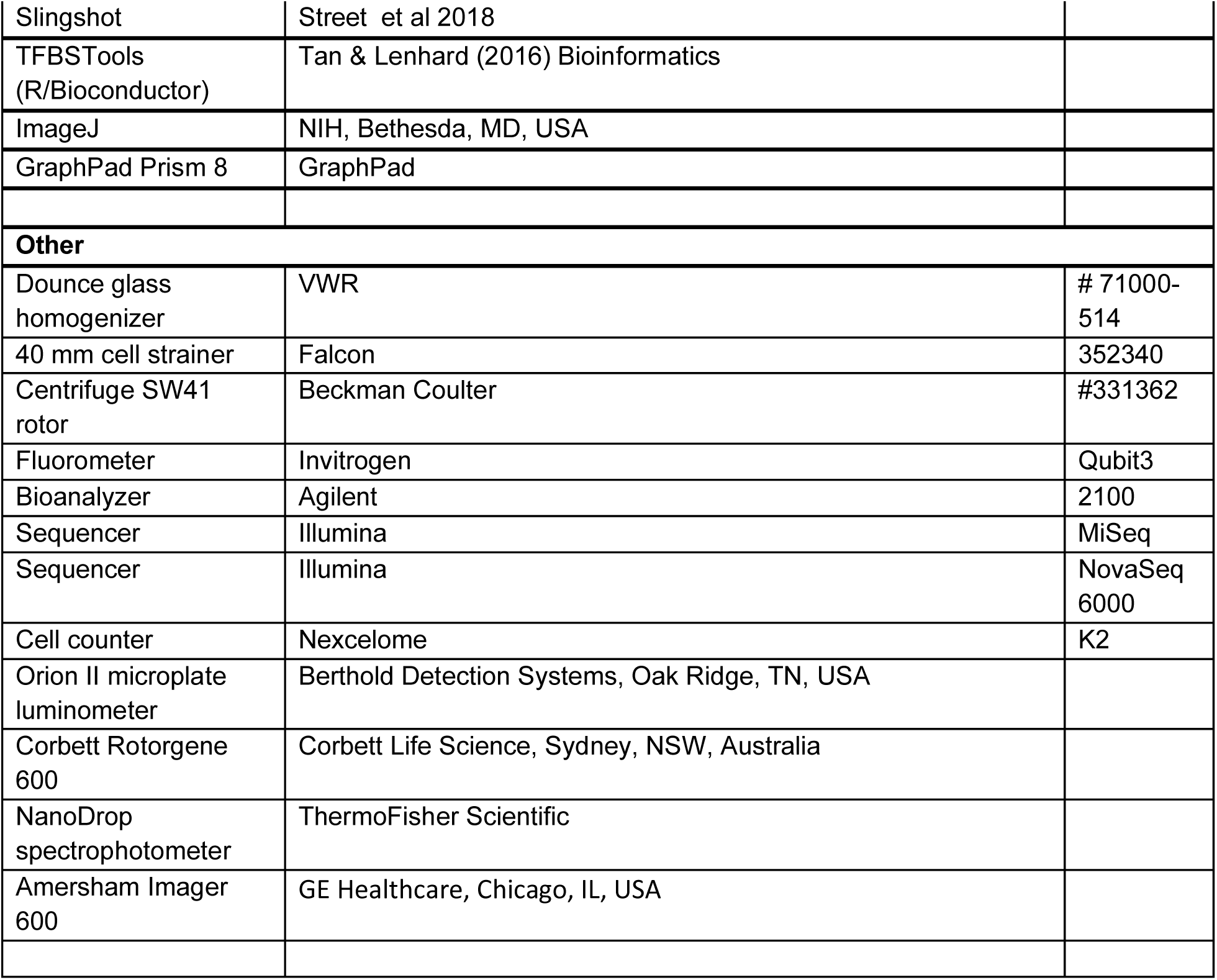

### Ethical compliance

We have complied with all ethical regulations and institutional protocols. All animal experiments were conducted following institutional and federal guidelines and were approved by the McGill University (Montreal, Quebec, Canada) and Goodman Cancer Institute DOW-A Animal Care Committee (Protocol 5204).

### Animals and estrous cycle assessment

Female mice (C57BL/6, strain number: 027) were purchased from Charles River Labs (Senneville, Quebec, Canada). To minimize mouse-handling stress, females were allowed to acclimate for one week before their estrous cyclicity was assessed at age 10 to 11 weeks for 1-2 weeks. For acclimation, the tail of the mice was massaged daily for 2 weeks before any blood collection from the tail tip. Each animal was individually handled using a paper roll, and 2 µL of blood was collected from the tail tip when the animal was in proestrus, as described in ^68^. Blood was collected either from the tail vein to confirm the LH surge at proestrus 6 pm (as assessed by vaginal cytology) using an in-house sandwich ELISA ^20^, or by cardiac puncture at all other stages of the cycle. Vaginal cytology was examined using 0.1% methylene blue, following established guidelines. Females were swabbed daily around 3 pm and were euthanized at 9 am on diestrus or at various times during proestrus (9 am, 6 pm, or 11 pm) or on the morning of estrus (2 am or 9 a) (refer to ^69^ Appendix 4). The animals had *ad libitum* access to food and water and were housed on a 14:10 lights on/lights off cycle.

Terminal blood collected by cardiac puncture was allowed to clot for 30 to 60 minutes at room temperature and then spun down at 3,000 rpm for 10 minutes to collect serum. Serum samples were stored at -80°C for hormone level determination. LH and FSH were measured using in-house ELISAs ^20^. Serum inhibins A and B were measured by commercial ELISAs (AnshLabs AL-161 and AL-163). Pituitary glands were dissected and snap-frozen until further analysis.

### Pituitary collection and nuclei isolation

Pituitaries were collected from C57BL/6 female mice aged 11 weeks. After staging of the mice, pituitaries were immediately snap-frozen following dissection, and stored at -80C until shipping for assay. The day of an assay and on ice, snap-frozen pituitaries were thawed and prepared based on a protocol described in ^11–14^. Briefly, RNAse inhibitor (NEB MO314L) was added to the homogenization buffer (0.32 M sucrose, 1 mM EDTA, 10 mM Tris-HCl, pH 7.4, 5mM CaCl2, 3mM Mg(Ac)2, 0.1% IGEPAL CA-630), 50% OptiPrep (Stock is 60% Media from Sigma; cat# D1556), 35% OptiPrep and 30% OptiPrep right before isolation. Each pituitary was homogenized in a dounce glass homogenizer (1ml, VWR cat# 71000-514), and the homogenate filtered through a 40 μm cell strainer. An equal volume of 50% OptiPrep was added, and the gradient centrifuged (SW41 rotor at 9200rpm; 4C; 25min). Nuclei were collected from the interphase, washed, resuspended either in 1X nuclei dilution buffer (10X Genomics), and counted (K2R Cellometer).

### Sc multiome assay

Sn multiome was performed following the Chromium Single Cell Multiome ATAC and Gene Expression Reagent Kits V1 User Guide (10x Genomics, Pleasanton, CA). Nuclei were counted (K2R Cellometer), transposition was performed in 10 ul at 37C for 60min targeting up to 10,000 nuclei, before loading of the Chromium Chip J (PN-2000264) for GEM generation and barcoding. Following post-GEM cleanup, libraries were pre-amplified by PCR, after which the sample was split into three parts: one part for generating the snRNAseq library, one part for the snATACseq library, and the rest was kept at -20C. SnATAC and snRNA libraries were indexed for multiplexing (Chromium i7 Sample Index N, Set A kit PN-3000262, and Chromium i7 Sample Index TT, Set A kit PN-3000431 respectively).

### Quality control (QC) and sequencing of libraries

QC and quantification of libraries were done by Bioanalyzer (High-Sensitivity DNA Bioanalyzer kit), Qubit fluorometer (Thermofisher), and mi-seq (Illumina). Sequencing was carried out at New York University Sequencing facility and at the New York Genome Center (NYGC) on an Illumina Novaseq using 98+26 paired-end reads.

### Sn data integration

#### Sn multiome primary analysis, dataset integration and cell type annotation

Cell Ranger ARC 1.0.0 (10x Genomics) was used to assign cellular barcodes and align the reads to the mouse reference genome (mm10) following 10x Genomics guidelines. After QC, subsequent analyses of the sn multiome data included cell type classification, identification of DEGs and DARs across cycle stages, pseudotime trajectory analysis, and inference of *cis*-regulatory circuitry and their biological implications.

#### snRNA primary

Ambient RNA removal was performed with decontX ^67^ by using only the filtered feature-barcode gene expression matrix for all samples except MF54. The resulting gene expression matrix and the filtered feature-barcode gene expression matrix per sample were read into a Seurat object ^70^. For MF54, which was a multiplexed dataset of two-samples (one male and one female), read counts of genes on the sex chromosomes were used to identify and retain female sample cells. The criteria used were: read counts for Xist > 0 and read counts of all genes on Y chromosome equal to 0. The barcodes of cells identified as female were also included in the snATAC analysis.

Cells with unusually high expression levels (nFeature >= 5000 and nCount >= 10000) were considered putative doublets and were removed from further analysis. Poor quality cells (nFeature <= 1000 or mitochondrial content >= 2%) were also filtered out.

#### snRNA integration

The merged Seurat dataset (visualized in a Uniform Manifold Approximation and Projection, UMAP) was examined for the presence of intrinsic batches. All the samples were then integrated following the Seurat integration pipeline using the ambient RNA corrected results while also correcting for the identified batches. Cell type annotation of the full dataset was performed using SingleR ^71^ with a 3-sample female mouse pituitary dataset serving as reference^12^.

#### snATAC primary

The fragment files from the Cell Ranger pipeline were processed and combined using ArchR [cite]. The addDoubletScores function was used for the initial doublet filtering. High quality cells with TSS enrichment >= 8, nucleosome ratio < 2, and doublet enrichment score <= 2 were selected for further analysis.

#### snATAC integration

Dimensionality reduction using the addIterativeLSI function was performed on the tile matrix based on the read counts in 500bp bins across the genome. UMAP was generated using the addUMAP function. To create a universal peak set for differential analysis, common peaks were called across subjects using the addReproduciblePeakSet function with the MACS2 peak caller ^72^.

### snRNA-snATAC joint processing

The cells that passed quality controls in both assay modalities were used in the joint snRNA and snATAC object and for downstream analysis. The snATAC ArchR object was exported to a Seurat/Signac object with adjustments (such as retaining the relevant cell barcodes) to fit the needs of sn-multiome data ^73^. The snRNA data matrices, dimensionality reduction results, and cell type annotation results were then combined with the snATAC Seurat object for downstream analysis.

### Trajectory analysis on snRNA

Additional filterings were done to ensure a consistent read count across samples for the downstream analysis in the gonadotrope subset (refer to sub-sections below). PCA and UMAP were redone, and trajectory and pseudotime analysis were done using slingshot with cluster by estrous cycle stages and “Proestrus 6pm” as the starting cluster ^23^.

### Cell type peak calling

For cell type-specific analysis of ATAC seq data, we used mac v2.1.0 to call ATAC seq peaks on all cells annotated to the same cell type, generating sets of cell type-specific peaks. All downstream analysis on chromatin accessibility peaks were conducted on these cell type-specific peak sets.

### Data cleaning in the gonadotrope population

Gonadotrope is the main cell type we wanted to focus on in this study, yet it is prone to doublets and ambient RNA contamination due to its small population and the presence of cell types with much larger populations, like somatotrope and lactotrope. We found that after standard doublet filtering and ambient RNA removal procedures conducted on all cells, many cells originally annotated as gonadotropes still presented somatotrope and lactotrope gene signatures, suggesting that there was still a high level of contamination in the gonadotrope population. Therefore, we conducted an additional round of manual doublet removal specifically for the gonadotrope population. Specifically, for each stage, we ran another round of clustering only on the gonadotropes of this stage. When we found more than one cluster, we then removed any clusters with high expression levels of Gh or Prl compared to the other clusters of this stage, as these clusters were likely doublets with somatotropes or lactotropes. All the results on gonadotrope presented in this manuscript were obtained after this procedure of data cleaning.

### Differential analysis of gene expression and chromatin accessibility

The differential analysis in this study mostly focused on the comparison between adjacent stages of the estrous cycle. The analysis of the three stages at 9am were conducted by comparison between adjacent 9am stages.

### Differentially expressed genes and differentially accessible regions

Differentially expressed genes (DEGs) were identified using the “FindMarkers” function from Seurat using “wilcox” test and a cutoff of Bonferroni corrected p values < 0.05 and absolute log2 fold change > 0.2. Differentially accessible regions (DARs) were identified using the “FindMarkers” function from Seurat using “LR” test and a cutoff of Bonferroni corrected p values < 0.05 and absolute log2 fold change > 0.2.

### Differential analysis on subsampled cells

To mitigate the influence of the number of cells on the number of identified DEGs and DARs when comparing across cell types and stages, we conducted an additional differential analysis with cell populations subsampled to the same sizes. Specifically, across all cell types and all stages, we found the cell type and stage that had the smallest number of cells (gonadotropes at estrus 9am, 278 cells). We then subsampled all cell types at all stages to the same size and ran differential analysis as described above. We subsampled 20 times and used the average numbers of DEGs and DARs across 20 runs as the final reported numbers. The exception to this procedure is the cell types at diestrus 9am, because we only collected 2 samples for this stage and we only had 94 gonadotropes at this stage. We treated diestrus 9am separately to avoid subsampling data from other stages to sizes that are too small. For diestrus 9am, we subsampled all other cell types of this stage to 94 cells and conducted differential analysis as described above. Therefore, the number of DEGs and DARs can be compared across cell types within diestrus 9am, but they were based on different numbers of cells compared to the results of all other stages. The comparison of the three 9am stages were also based on the same subsampling strategy.

### Gene dynamics analysis

Gene expression temporal trajectories at pseudobulk level were clustered to reveal different patterns of gene expression across the estrous cycle. Specifically, for each cell type we calculated the gene expressions at sample level (pseudobulk level) and then averaged at stage level so that the stage-level pseudobulk expressions were not dominated by samples with larger number of cells. Each gene was then represented by a vector of its temporal expression values at each of the six stages. Genes were clustered using k means clustering on these stage level expression values. Only differential genes with significant expression change reported in Figure 2A were included in this analysis.

The number of clusters in the kmeans clustering were determined by both the “fviz_nbclust” function from the “factoextra” using gap statistic, as well as visual inspection of the similarities between clusters under different choices of k. We chose k=7 for gonadotropes and k=6 for lactotropes.

### Gene regulatory circuitry analysis

We used MAGICAL to analyze the GRCs ^16^. Each circuit is composed of a transcription factors (TF), a target gene, and a cis-regulatory region the TF interacts with to regulate the expression of the target gene. MAGICAL was originally developed for cell type-level pseudobulk data. To utilize the RNAseq and ATACseq data matched at the sc level, we aggregated single cells to meta cells and use the pseudobulk data at meta cell-level as the input for MAGICAL.

#### Metacells

We used SEACells ^26^ with default parameters to construct meta cell from single cell data. Specifically, meta cell construction was done using the RNAseq data and the PCA reduction as the low dimensional kernel as the inputs to SEACells. For gonadotropes, we chose to construct 50 meta cells in total based on the recommendation of targeting 1 meta cell per 50 single cells from SEACells documentation. For lactotropes, due to the large quantity of lactotrope cells and run time limitations, we subsampled 2000 lactotropes from each sample and then targeted 1 meta cell per 250 single cells, resulting in 72 meta cells.

#### MAGICAL

The analysis using MAGICAL ^16^ focused on identifying the regulatory circuitry of the DEGs reported in Figure 2A. For the candidate cis-regulatory regions, we chose chromatin peaks within a 200kb window centered around the TSS. We also filter for chromatin peaks that were differentially accessible between any of the two adjacent stages, so that the regulatory circuits we constructed were relevant to the molecular changes in estrous cycle. Specifically, in gonadotropes, due to the lack of significantly differential peaks under the default cutoff, we used a relaxed cutoff of FDR < 0.005 to select differential peaks to use in the MAGICAL analysis. MAGICAL uses correlated patterns between chromatin accessibility and gene expression across meta cells to construct circuits and doesn’t rely on very strict differential signal calling. In lactotropes, we used the default cutoff for differential peaks as reported in Figure 2B because there were already abundant differential peaks as candidate peaks for MAGICAL analysis.

#### TF module activity

After circuit construction, we aggregated the circuits by the regulatory TFs. A TF module is defined as all cis-regulatory regions and target genes regulated by a common TF. For a specific TF X that interacts with cis-regulatory region (chromatin peak) Y and regulate the expression of gene Z, we define the “target circuit activity” as:

Target_circuit_activity (peak Y, gene Z) = Accessibility (peak Y) * Expression (Gene Z) The activities of each target circuit of each TF were calculated using sample-level pseudobulk peak accessibility and gene expression values.

The TF module activity in each sample was calculated by aggregating the activities of all the target circuits in this sample under this TF. Specifically, we used the “AUCell_run” function from the “AUCell” package to aggregate the activities of multiple target circuits into a single score for each TF in each sample. The function uses an AUC score to measure the rank of the activities of these target circuits over all target circuits in each sample. The result is a TF-by-sample matrix with TF module activity scores of each TF in each sample. Finally, the sample-level TF module activities were average to stage-level TF module activities.

#### Regulatory network analysis of the gene dynamics clusters

For each cluster of gene expression dynamics, we found the TFs with the largest number of target genes within this cluster. The resulting regulatory networks connecting TFs to target genes were visualized using the Gephi software.

### DNA constructs

The wild-type -1990/+1 murine Fshb-luc promoter-reporter ^35^, full-length wildtype Enh3 and Enh4 in *Fshb*-luc promoter reporter were described previously ^31^. HA-human TGFBR1 (T204D, ALK5TD) was provided by Dr. Peter Scheiffele (Columbia University).

Human ETS2 WT expression vector was obtained from Addgene (Watertown, Massachusetts, USA, RRID:Addgene_28128) and optimized for overexpression by mutagenesis.

Mutagenic primers (Table S1) were designed as complementary oligonucleotides with the desired substitutions centered within the primer sequence. Briefly, whole-plasmid amplification was performed using Pfu-Turbo (Agilent Technologies, Santa Clara, CA, USA; Cat. No. 600250) and the parental reporter plasmid as template, followed by DpnI (New England Biolabs, Ipswich, MA, USA; Cat. No. R0176) digestion to remove methylated/hemimethylated parental DNA. The digested reactions were transformed into chemically competent *E. coli*, and individual colonies were screened by plasmid miniprep prior to sequence verification. All mutant constructs were confirmed by Sanger sequencing (Genome Quebec). All mutant promoter/enhancer reporters were generated using the QuikChange protocol and primers in **Table S11**.

### Predicted effect of ETS2 protein and motif mutation on ETS2 binding

#### ETS2 protein mutation effect

mCSM-NA (https://biosig.lab.uq.edu.au/mcsm_na/prediction) ^74^ was used to predict the R419D mutation effect on the binding affinity of ETS2. The ETS2-DNA complex structure was downloaded from PDB (ID: 4BQA) and used as the input for mCSM-NA. The predicted effect of R419D mutation is reduced affinity with ΔΔG=1.265 kcal/mol, which corresponds to a 7-fold change of the dissociation constant Kd when T=310K.

#### ETS2 DNA motif mutation effect

The exact location of the binding site of Ets2 within Enh4 was identified by MAGICAL to be chr2:107127400-107127412. The wildtype DNA sequence is GAAGGGAAGGAAA and the mutated DNA sequence is GAAGGGAATTAAA. To predict the binding probability change from the mutations on the DNA sequence, we used the position frequency matrix (PFM) of ETS2 downloaded from HOCOMOCO v14 (motif ID: ETS2.H14CORE.1.P.B) ^75^. The mutation from GG to TT is predicted to cause a 5-fold decrease of binding probability according to the PFM.

### Cell line

Immortalized murine gonadotrope-like LβT2 (LβT2b cells (RRID:CVCL 0398) provided by Dr. Pamela Mellon (University of California, San Diego, CA, USA) were cultured in DMEM with 10% (v/v) fetal bovine serum ^76^. All cells were cultured at 37°C with 5% CO_2_ in a humidified incubator.

### Promoter-reporter assays

Promoter-reporter assays were performed as previously described ^77^. Briefly, LβT2b cells ^76^ were seeded at a density of 150,000 cells per well in 48-well plates. The next day, the cells were transfected with 225 ng/well of the indicated reporter plasmid constructs, with or without 5 ng pcDNA3.0, ETS2 WT, or ETS2 R419D using Lipofectamine 3000 (L3000015, ThermoFisher Scientific, Burlington, ON, Canada) following the manufacturer’s protocol. Twenty-four hours after transfection, cells were serum-starved overnight. The following day, cells were treated with activin A (0.05 or 0.5 nM; 388-AC-050, R&D Systems, Minneapolis, MN, USA) for 6 h, or with serum-free medium used as the vehicle control. Cells were then lysed in 50 µL/well passive lysis buffer (25 mM Tris-phosphate [pH 7.8], 10% [v/v] glycerol, 1% [v/v] Triton X-100, 1 mg/mL bovine serum albumin, 2 mM EDTA) for 10 min at room temperature with agitation. Lysates were clarified, and 20 µL of the supernatant was combined with 100 µL assay buffer (15 mM potassium phosphate [pH 7.8], 25 mM glycylglycine, 15 mM MgSO₄, 4 mM EDTA, 2 mM ATP, 1 mM DTT, 0.04 mM D-luciferin). Luciferase activity was measured using an Orion II microplate luminometer (Berthold Detection Systems, Oak Ridge, TN, USA). Assays were performed in technical duplicates or triplicates, and each experiment was repeated at least three times as indicated in the corresponding figure legends.

### Co-expression of ETS2 and ALK5TD

LβT2 cells were seeded in 6-well plates at 1,500,000 cells/well and immediately transfected using FuGENE® 4K Transfection Reagent (Promega, Madison, WI, USA; Cat. #E5911) via “fast-forward” transfection before the cells adhere to plates. Transfection master mixes were prepared in serum-free DMEM by first mixing plasmid DNA with DMEM, then adding FuGENE 4K at a 3:1 (µL FuGENE:µg DNA) ratio, followed by immediate mixing, and incubation at room temperature for 15 min. A total of 100 µL transfection master mix was added per well, and cells were incubated overnight. For each well, 500 ng/well of ETS2 WT or ETS2R419D plasmid was transfected with or without 500 ng/well constitutively active ALK5 (ALK5TD), with total DNA normalized using pcDNA3.0.

The next day, cells were serum-starved, and the following day cells were lysed in TRIzol for RNA isolation. In parallel, a separate 6-well plate was transfected side-by-side using the same transfection mixes and harvested for immunoblotting to verify expression of transfected constructs.

### Reverse transcription and quantitative PCR

Total RNA was isolated from homogenized pituitaries using TRIzol (15596018, Invitrogen, Waltham, MA, USA) according to the manufacturer’s instructions, and RNA concentration was measured by NanoDrop spectrophotometry. For all experiments, 200 ng total RNA was treated with RQ1 DNase and reverse-transcribed using random hexamer primers (C1181, Promega) and Moloney murine leukemia virus reverse transcriptase (M1701, Promega). Resulting cDNA was diluted 1:2 in DEPC-treated H₂O before qPCR. Quantitative PCR was performed with EvaGreen (ABMMmix, Diamed, Mississauga, ON, Canada) using primers listed in **Supplementary Table S11** on a Corbett Rotorgene 600 (Corbett Life Science, Sydney, NSW, Australia). Primer efficiency and specificity were validated before use. Relative transcript abundance was calculated using the 2^-ΔΔCt^ method and normalized to *Gapdh*.

### Western blotting

For each sample, 25 µg total protein was mixed with 5× Laemmli sample buffer (final 1×; 250 mM Tris-HCl pH 6.8, 10% SDS, 50% glycerol, 0.02% bromophenol blue, 10% β-mercaptoethanol) and heated to denature proteins. Proteins were separated by SDS–PAGE on a 10% resolving gel prepared from a 30% (w/w) acrylamide/bis-acrylamide (29:1) solution in running buffer (25 mM Tris, 250 mM glycine, 0.1% SDS, pH 8.3). Proteins were transferred to Protran nitrocellulose membranes (GE 10600001; MilliporeSigma, Oakville, Ontario, Canada) in Towbin transfer buffer (25 mM Tris, 192 mM glycine, pH 8.3, 20% methanol). Membranes were blocked in 5% (w/v) milk in TBST (TBS: 150 mM NaCl, 10 mM Tris-HCl pH 8.0, supplemented with 0.05% (v/v) Tween-20) and incubated overnight at 4°C with agitation with an anti-FLAG primary antibody diluted in blocking buffer (1:1000, Sigma-Aldrich Cat# F7425; AB_439687) for ETS2. The next day, membranes were washed in TBST and incubated with HRP-conjugated anti-rabbit secondary antibody (1:5000; AP182P; MilliporeSigma; RRID:AB_92591) in blocking buffer for 1 h at room temperature with agitation. For detecting ALK5TD, anti-HA primary antibody (1:1000, Sigma-Aldrich Cat#H9658; RRID:AB_260092) and HRP-conjugated anti-mouse secondary antibody (1:5000; BioRad 1706516; RRID:AB_2921252) were used. After additional TBST washes, bands were detected using enhanced chemiluminescence substrate (NEL105001, PerkinElmer, Waltham, MA, USA) and imaged on an Amersham Imager 600 (GE Healthcare, Chicago, IL, USA).

Band intensities were quantified in ImageJ (NIH, Bethesda, MD, USA) using the Gel Analysis tool. Within each experiment, signals were normalized to the ETS2 or ALK5TD only condition, which was set to 1.

### Statistical analyses

Luciferase assay data in LβT2 were log-transformed before analysis by two-way ANOVA, followed by Holm-Šidák test. Western blots were analyzed using one-way ANOVA followed by Tukey’s post hoc tests. Statistical analyses were performed using Prism 8, GraphPad. Alpha was set to P < 0.05.

## Supplementary tables

### Supplementary tables (large) provided as individual spreadsheets

**Table S5.** Full list of DEGs identified between two consecutive estrous cycle time points in each pituitary cell type

**Table S6.** Full list of DARs identified between two consecutive estrous cycle time points in each pituitary cell type

**Table S7.** List of the genes in each of the 6 temporal dynamics clusters identified in the lactotropes across estrous cycle time points. Related to Fig. 4A.

**Table S8.** List of the genes in each of the 7 temporal dynamics clusters identified in the gonadotropes across estrous cycle time points. Related to Fig. 4B.

**Table S9.** Full list of lactotrope GRCs identified using the MAGICAL framework

**Table S10.** Full list of gonadotrope GRCs identified using the MAGICAL framework

**Table S11:** Primers used in mutagenesis and qPCR experiments

