## Supplemental data for "Single-nucleus multiomic analysis reveals modulation of the gene regulatory circuit landscape in pituitary cell types during mouse estrous cycle"

* Authors contributed equally to this work.

+ Co-corresponding authors share senior authorship.

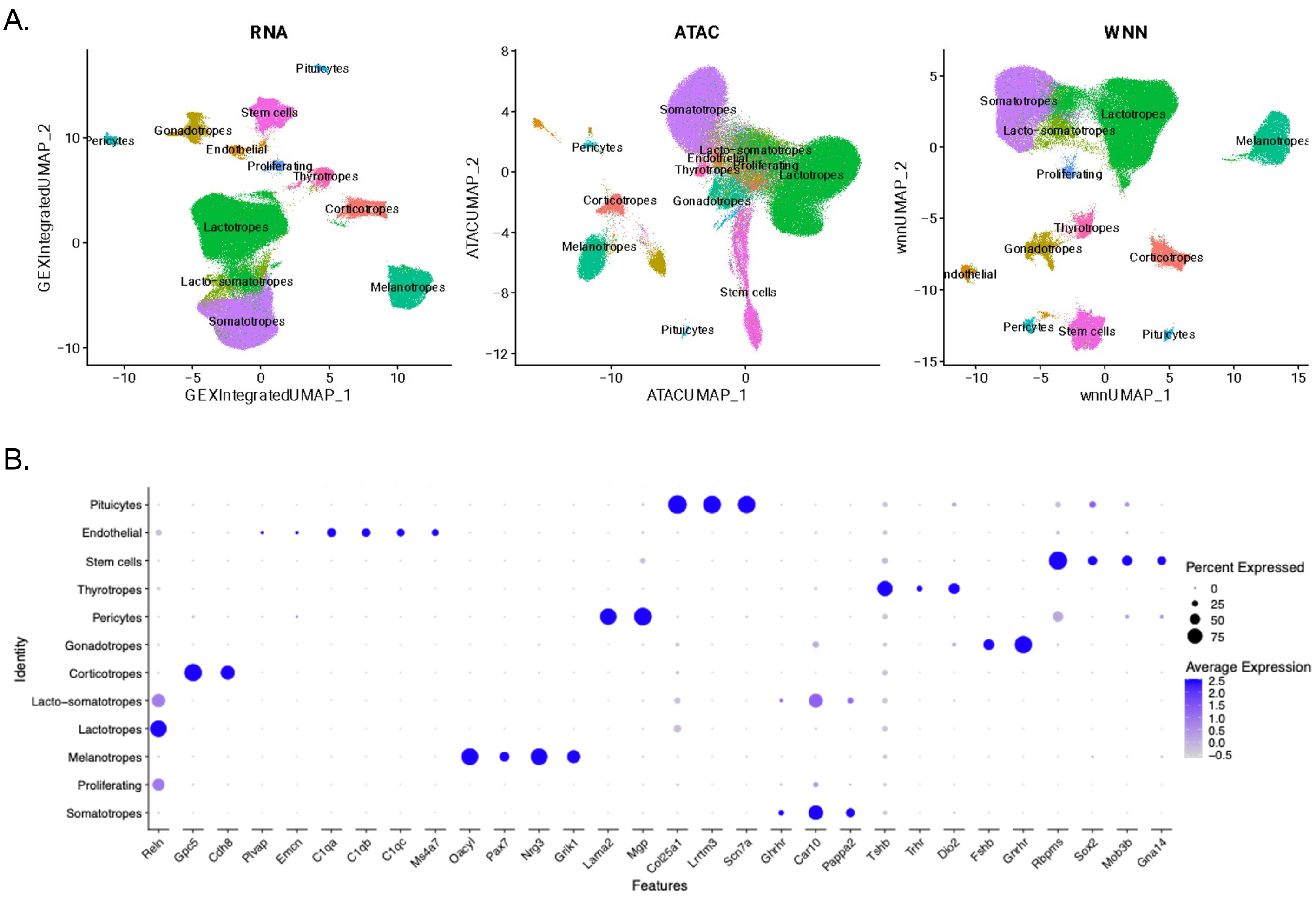
A.
RNA
ATAC
WNN
GEXIntegratedUMAP_1
GEXIntegratedUMAP_2
ATACUMAP_1
ATACUMAP_2
wnnUMAP_1
wnnUMAP_2
Pituicytes
Stem cells
Gonadotropes
Pericytes
Endothelial
Proliferating
Thyrotropes
Corticotropes
Lactotropes
Lacto-somatotropes
Melanotropes
Somatotropes
B.
Identity
Features
Percent Expressed
Average Expression
Pituicytes
Endothelial
Stem cells
Thyrotropes
Pericytes
Gonadotropes
Corticotropes
Lacto-somatotropes
Lactotropes
Melanotropes
Proliferating
Somatotropes
Reln
Gpc5
Cdh8
Plvap
Emcn
C1qa
C1qb
C1qc
Ms4a7
Oacyl
Pax7
Nrg3
Grik1
Lama2
Mgp
Col25a1
Lrrtm3
Scn7a
Ghrhr
Car10
Pappa2
Tshb
Trhr
Dio2
Fshb
Gnrhr
Rbpms
Sox2
Mob3b
Gna14

Fig S1

**Fig. S1: Pituitary cell type identification by sn analysis. A.** UMAPs of the 18 murine pituitaries across estrous cycles showing all identified cell types. Showing the individual modalities, snRNAseq (*Left panel*), snATACseq (*Middle panel*) and the integrated snRNA/snATAC seq (VWN, *Right panel*). The cell types are color-coded. **B.** Dot plot correlating established markers used for cell type annotation in **A**. with their cell type. Shown are the percentages of cells expressing the indicated markers, along with the average level of expression of each marker within those cells. Refer to Figure 1.

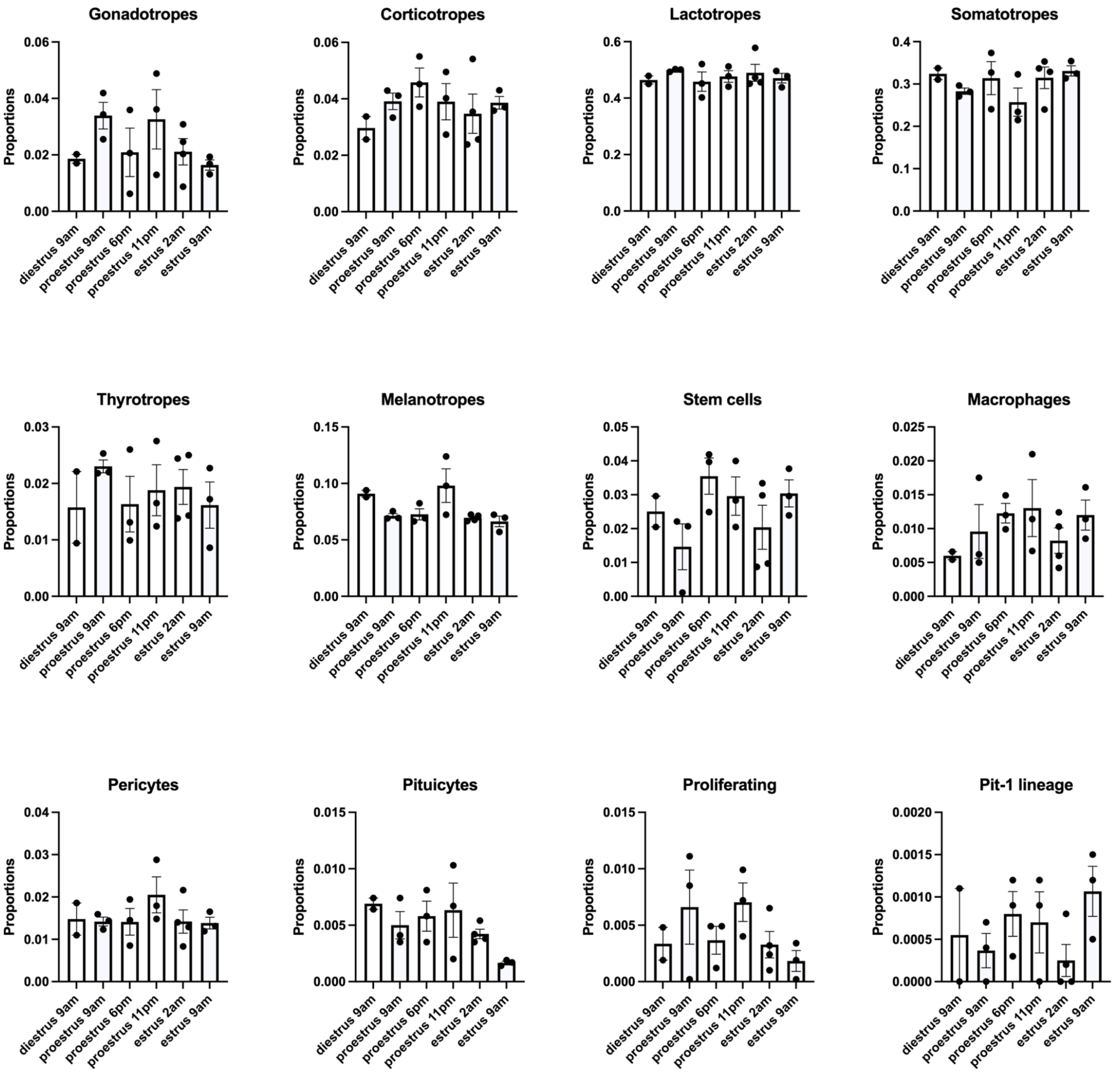
Gonadotropes
Proportions
0.06
0.04
0.02
0.00
Corticotropes
Proportions
0.06
0.04
0.02
0.00
Lactotropes
Proportions
0.6
0.4
0.2
0.0
Somatotropes
Proportions
0.4
0.3
0.2
0.1
0.0
Thyrotropes
Proportions
0.03
0.02
0.01
0.00
Melanotropes
Proportions
0.15
0.10
0.05
0.00
Stem cells
Proportions
0.05
0.04
0.03
0.02
0.01
0.00
Macrophages
Proportions
0.025
0.020
0.015
0.010
0.005
0.000
Pericytes
Proportions
0.04
0.03
0.02
0.01
0.00
Pituicytes
Proportions
0.015
0.010
0.005
0.000
Proliferating
Proportions
0.015
0.010
0.005
0.000
Pit-1 lineage
Proportions
0.0020
0.0015
0.0010
0.0005
0.0000
diestrus 9am
proestrus 9am
proestrus 6pm
proestrus 11pm
estrus 2am
estrus 9am

Fig S2

**Fig. S2: Cell type proportions across the estrous cycle, identified from the snRNAseq data.** Each bar graph correspond to an individual cell type, with each bar corresponding to the indicated estrous cycle stage. Related to **Fig. 1**.

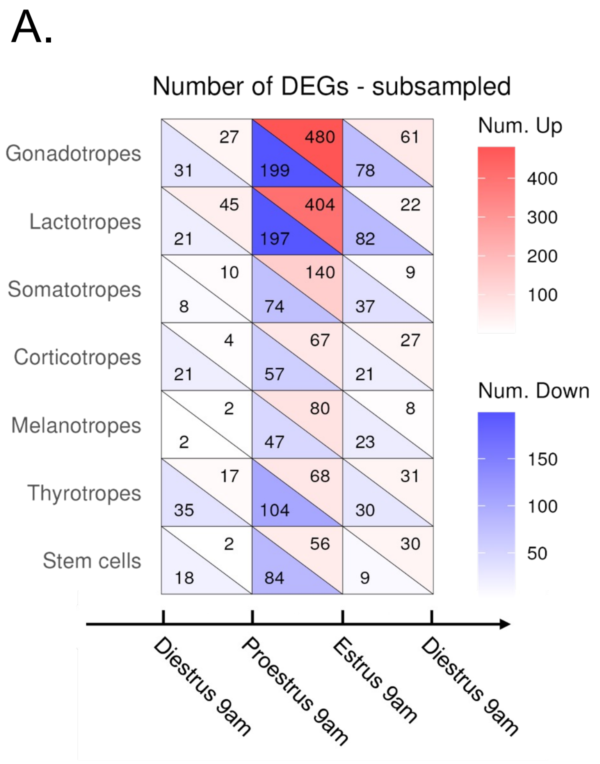
A.
Number of DEGs - subsampled
Gonadotropes
Lactotropes
Somatotropes
Corticotropes
Melanotropes
Thyrotropes
Stem cells
27
31
480
199
61
78
45
21
404
197
22
82
10
8
140
74
9
37
4
21
67
57
27
21
2
2
80
47
8
23
17
35
68
104
31
30
2
18
56
84
30
9
Num. Up
400
300
200
100
Num. Down
150
100
50
Diestrus 9am
Proestrus 9am
Estrus 9am
Diestrus 9am
B.
Number of DARs - subsampled
Gonadotropes
Lactotropes
Somatotropes
Corticotropes
Melanotropes
Thyrotropes
Stem cells
0
0
20
36
1
0
0
0
95
78
0
0
0
1
10
10
0
0
0
0
8
0
0
0
0
0
11
3
0
2
0
1
1
2
1
0
0
0
3
0
0
0
Num. Down
60
40
20
0
Num. Up
75
50
25
0
Diestrus 9am
Proestrus 9am
Estrus 9am
Diestrus 9am

Fig S3

**Figure S3. Analysis of circadian influence on DEGs and DARs.** Number of DEGs (**A**) and DARS (**B**) per cell type between estrous cycle stages studied at 9 AM, with the number of cells per cell type being subsampled across cell types and time points, at the exception of Diestrus 9 am samples where the number of cells were subsampled across cell types only because only two samples were assayed at that stage. Upregulated genes are in red and downregulated genes are in blue; regions of increased accessibility in red and regions of decreased accessibility in blue. Related to **Fig. 2**.

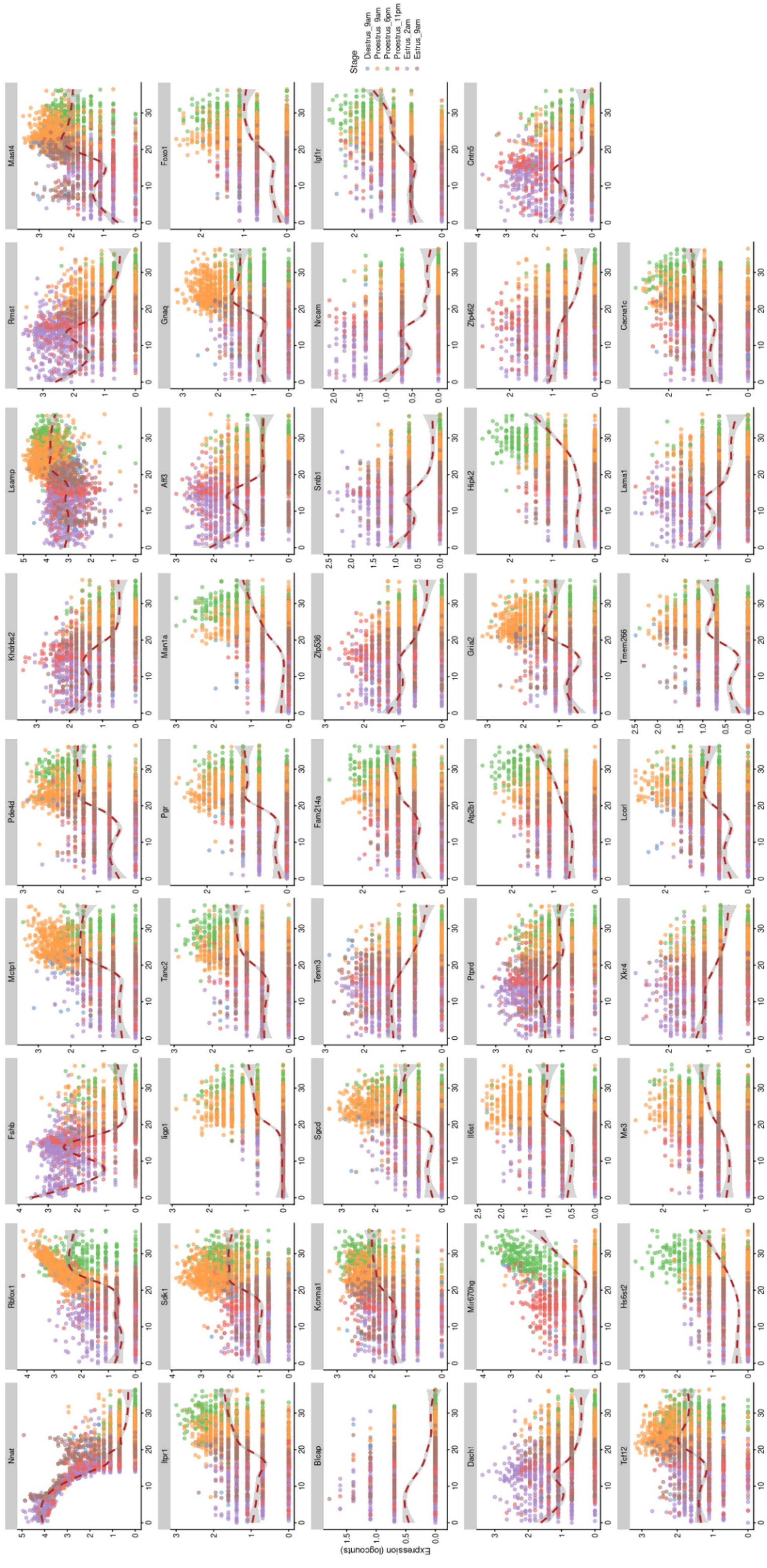

Fig S4

**Figure S4. Representation of genes contributing to pseudotime.** Represented are the 44 top genes contributing to pseudotime trajectories. Each dot represent a cell. The color code indicates the estrous cycle stage. The dotted line represents the median pseudotime. Related to **Fig. 3**.

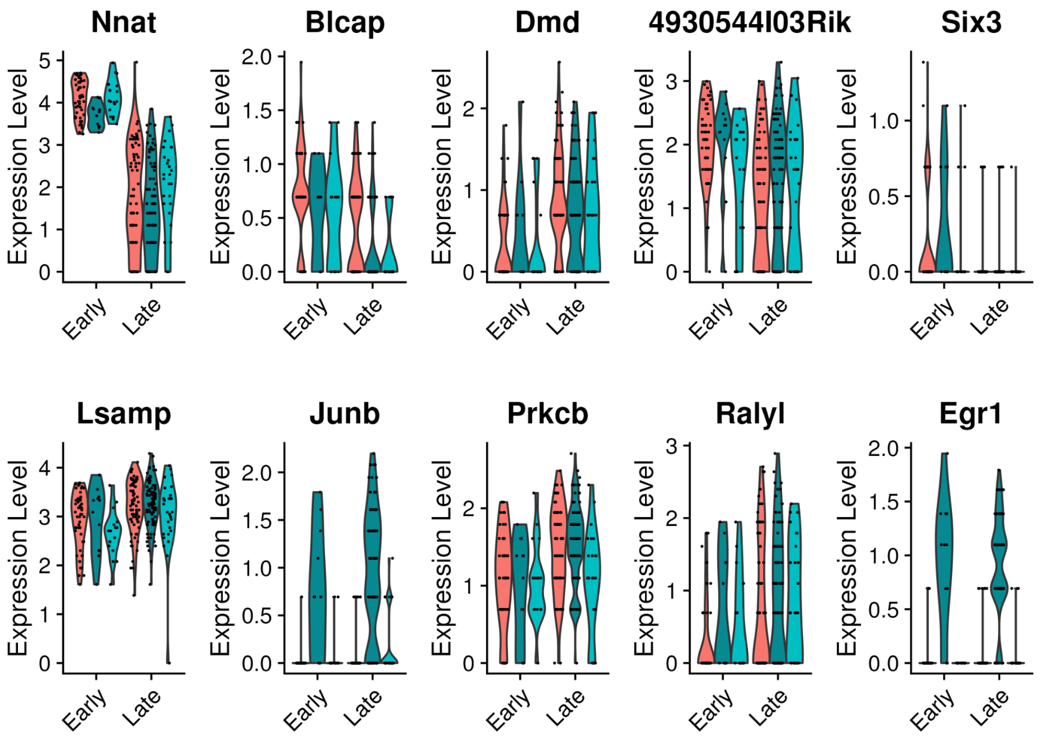
Nnat
Blcap
Dmd
4930544I03Rik
Six3
Lsamp
Junb
Prkcb
Ralyl
Egr1
Expression Level
Early
Late

Fig S5

**Figure S5. Violin plots of transcript expression for the indicated genes in the two Estrus 9 am pseudotime clusters of gonadotropes.** Related to **Fig. 3**.

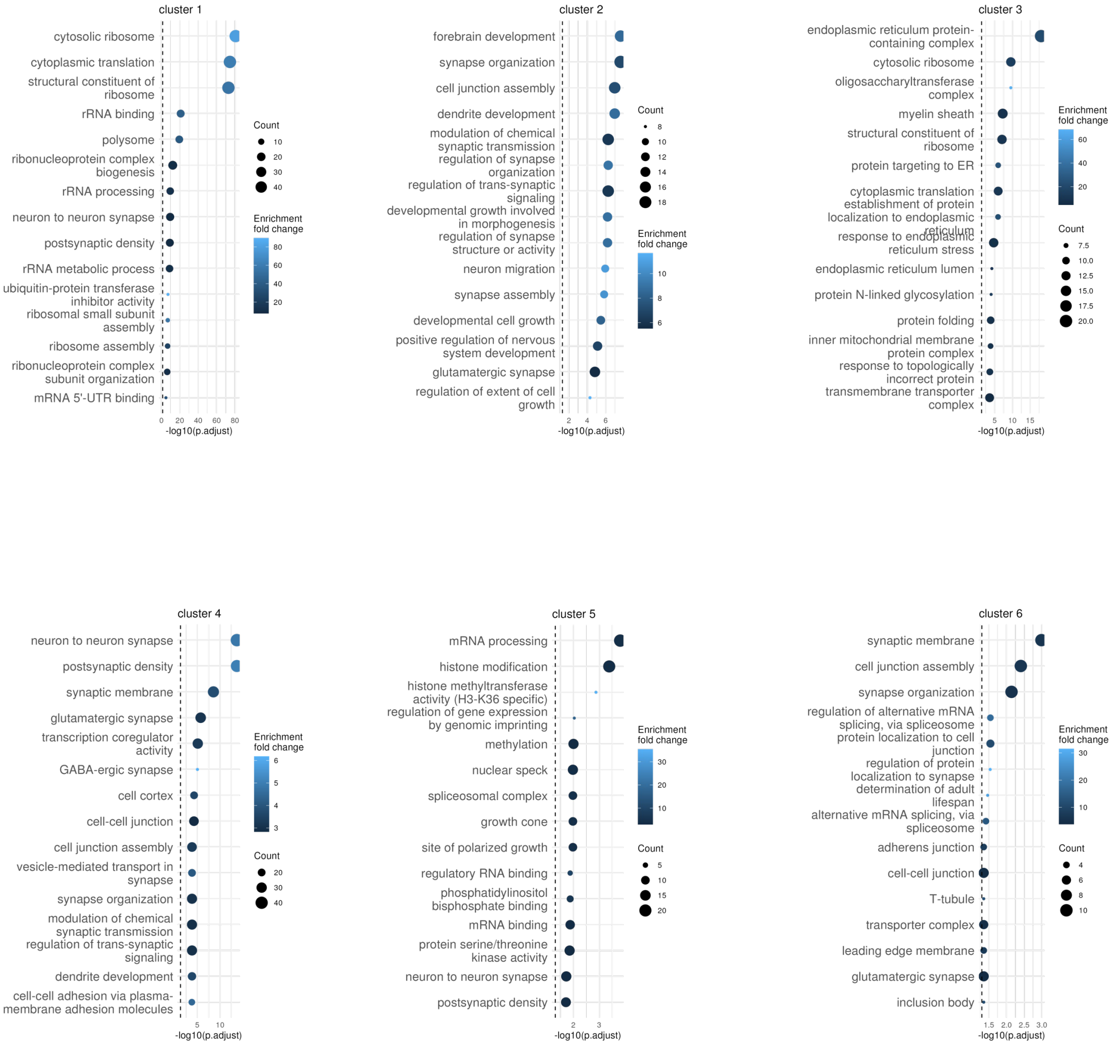

Lactotropes

Fig S6

**Figure S6. Pathway enrichment analysis on the temporal trajectory gene clusters identified in the lactotropes.** Shown are the biological pathways that are over-represented in each of the gene clusters depicted in **Fig. 4A**. Related to **Fig. 4**.

A.

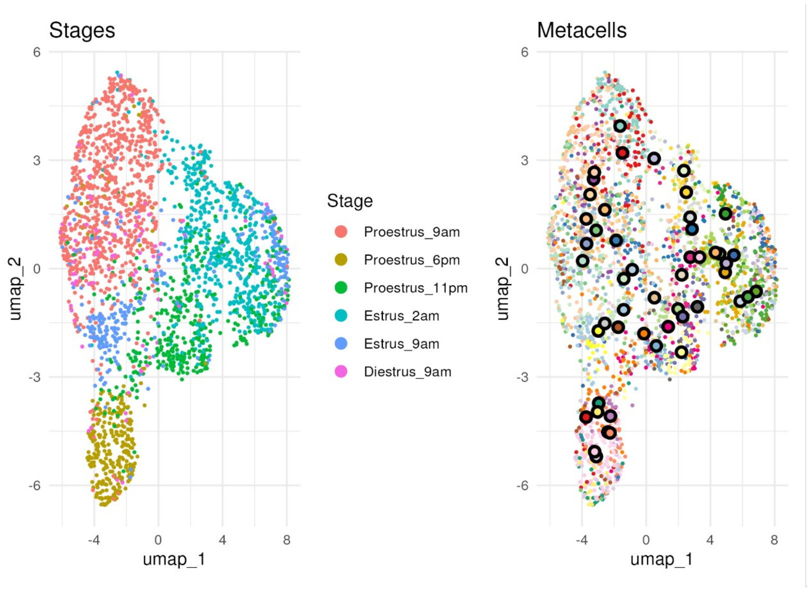
Stages
Metacells
Stage
Proestrus_9am
Proestrus_6pm
Proestrus_11pm
Estrus_2am
Estrus_9am
Diestrus_9am
umap_1
umap_2
6
3
0
-3
-6
-4
0
4
8

Fig S7

**Figure S7. Grouping of all gonadotrope cells into metacells.** *Left*, snRNA-seq UMAP representation of the gonadotrope cell cluster, with labeling by estrous cycle stage/time assignment. Estrous cycle stages are color-coded as indicated. *Right*, Gonadotrope cells from *Left* were grouped into metacells by applying the SEACells method (Persad 2023 https://pmc.ncbi.nlm.nih.gov/articles/PMC10713451/; see Methods) to the snRNAseq data. Related to **Fig. 5**.

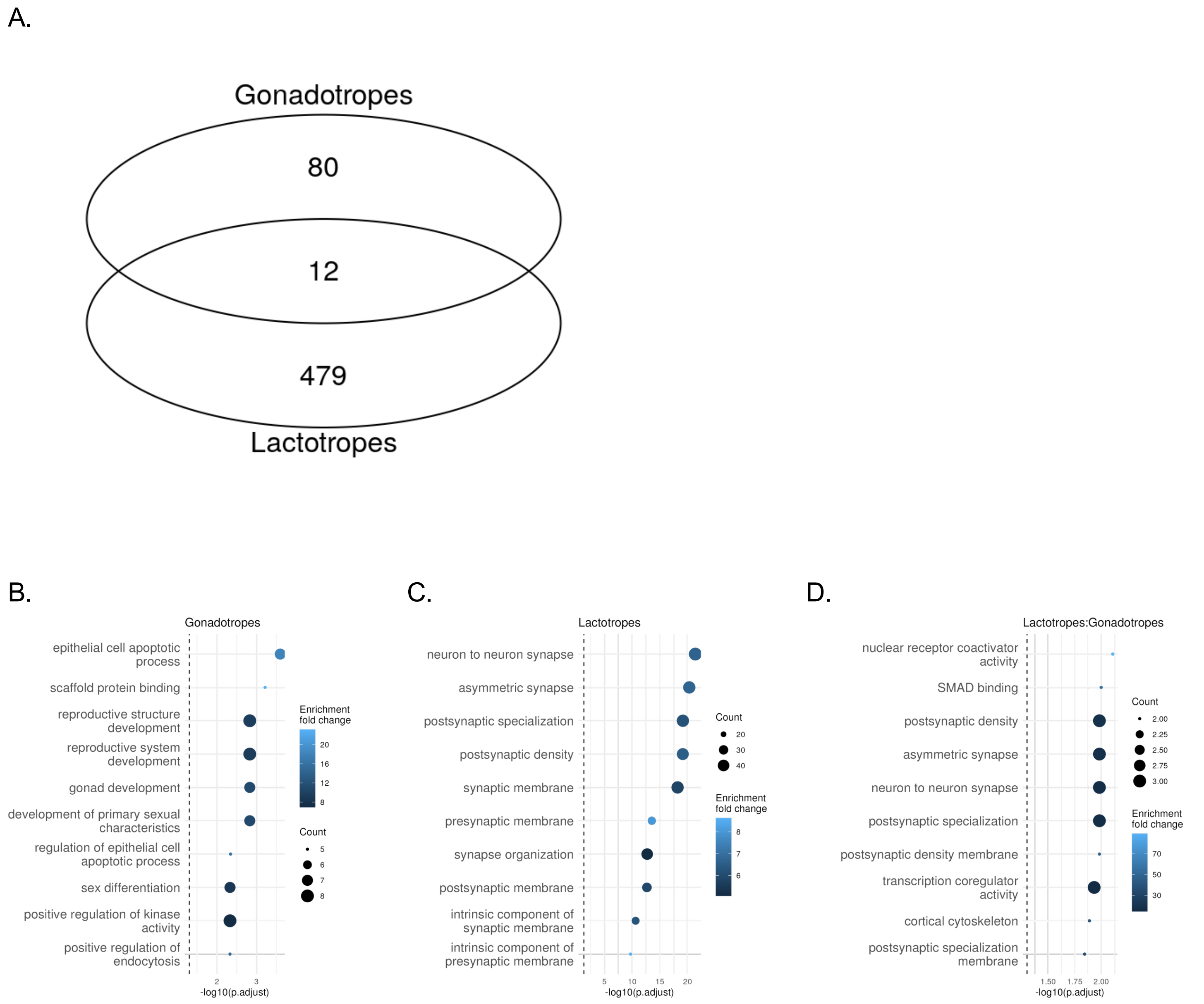
A.
Gonadotropes
80
12
479
Lactotropes
B.
Gonadotropes
epithelial cell apoptotic process
scaffold protein binding
reproductive structure development
reproductive system development
gonad development
development of primary sexual characteristics
regulation of epithelial cell apoptotic process
sex differentiation
positive regulation of kinase activity
positive regulation of endocytosis
Enrichment fold change
Count
-log10(p.adjust)
C.
Lactotropes
neuron to neuron synapse
asymmetric synapse
postsynaptic specialization
postsynaptic density
synaptic membrane
presynaptic membrane
synapse organization
postsynaptic membrane
intrinsic component of synaptic membrane
intrinsic component of presynaptic membrane
Count
Enrichment fold change
-log10(p.adjust)
D.
Lactotropes:Gonadotropes
nuclear receptor coactivator activity
SMAD binding
postsynaptic density
asymmetric synapse
neuron to neuron synapse
postsynaptic specialization
postsynaptic density membrane
transcription coregulator activity
cortical cytoskeleton
postsynaptic specialization membrane
Count
Enrichment fold change
-log10(p.adjust)

Fig S8

**Figure S8. The gonadotrope- and the lactotrope-specific Smad3 modules share target genes. A.** Shown is the target gene overlap between the gonadotrope- and the lactotrope-specific Smad3 modules. **B-D.** Biological pathways that are over-represented among the gonadotrope-specific target genes (**B**), the lactotrope-specific target genes (**C**), and the target genes shared between gonadotropes and lactotropes (**D**). Related to **Fig. 5**.

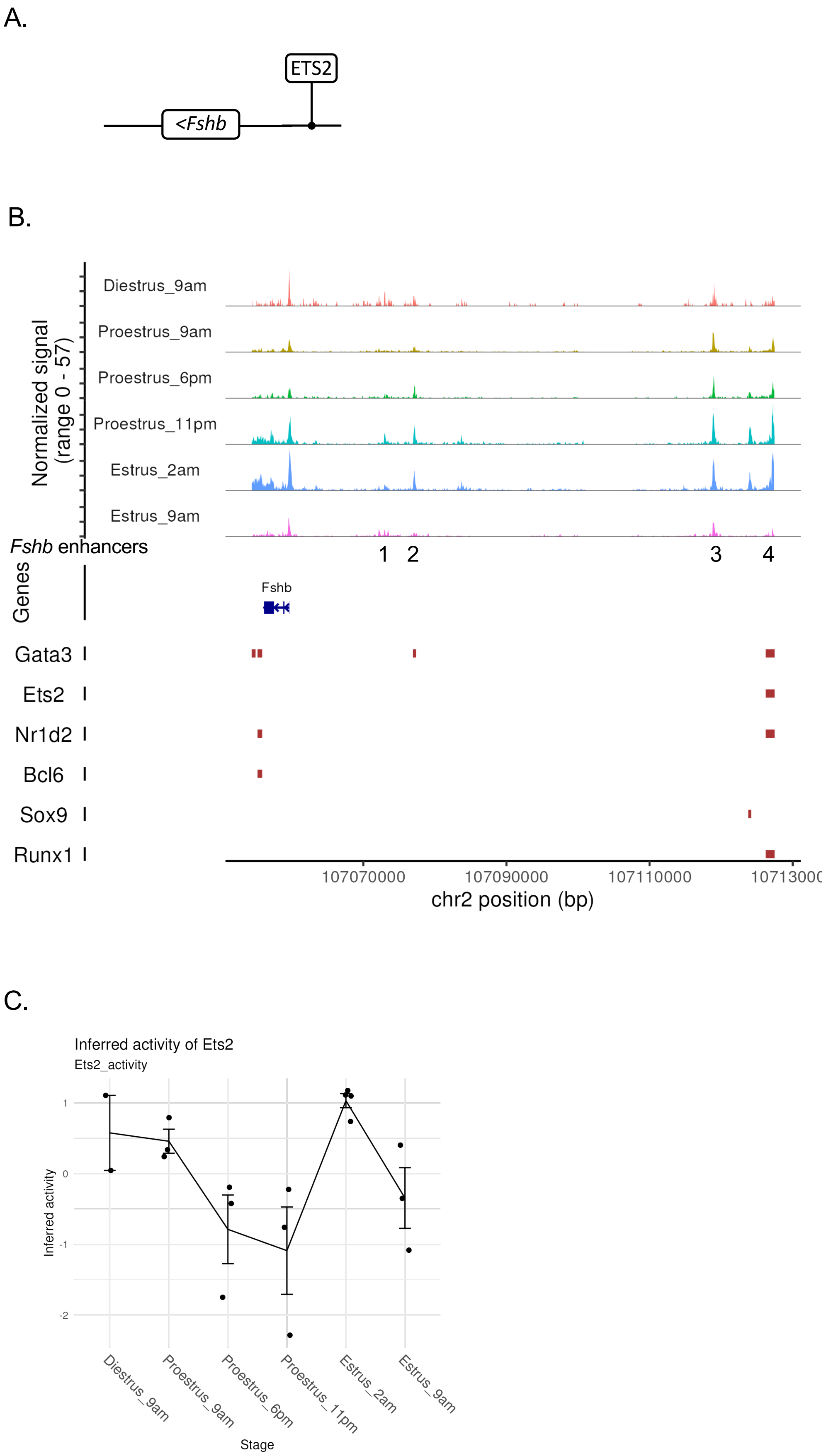
A.
ETS2
<Fshb
B.
Normalized signal (range 0 - 57)
Diestrus_9am
Proestrus_9am
Proestrus_6pm
Proestrus_11pm
Estrus_2am
Estrus_9am
Fshb enhancers
1 2 3 4
Genes
Fshb
Gata3
Ets2
Nr1d2
Bcl6
Sox9
Runx1
107070000
107090000
107110000
chr2 position (bp)
C.
Inferred activity of Ets2
Ets2_activity
Inferred activity
1
0
-1
-2
Diestrus_9am
Proestrus_9am
Proestrus_6pm
Proestrus_11pm
Estrus_2am
Estrus_9am
Stage

Fig S9

**Figure S9. Chromatin accessibility upstream of the *Fshb* gene and top regulatory TFs. A.** Schematic of the ETS2-*Fshb* GRC. **B.** Shown are the chromatin accessibility tracks of the *Fshb* gene and 5’ flanking region at each of the estrous cycle time points. The positions of the *Fshb* enhancer domains Enh1-Enh4 are signified with numbers (1-4). The positions of the indicated TF binding sites are denoted with red rectangles. Presented at the bottom of the figure is a schematic view of the genomic region holding the *Fshb* gene and 5’ flanking region, with genomic coordinates. The precise genomic coordinates of the enhancer domains Enh1-Enh4 are provided at the top. **C.** Line graph illustrating the TF activity of ETS2 in the ETS2-*Fshb* GRC in the gonadotropes across the estrous cycle, as inferred from MAGICAL. Related to **Figs. 5 and 6**.

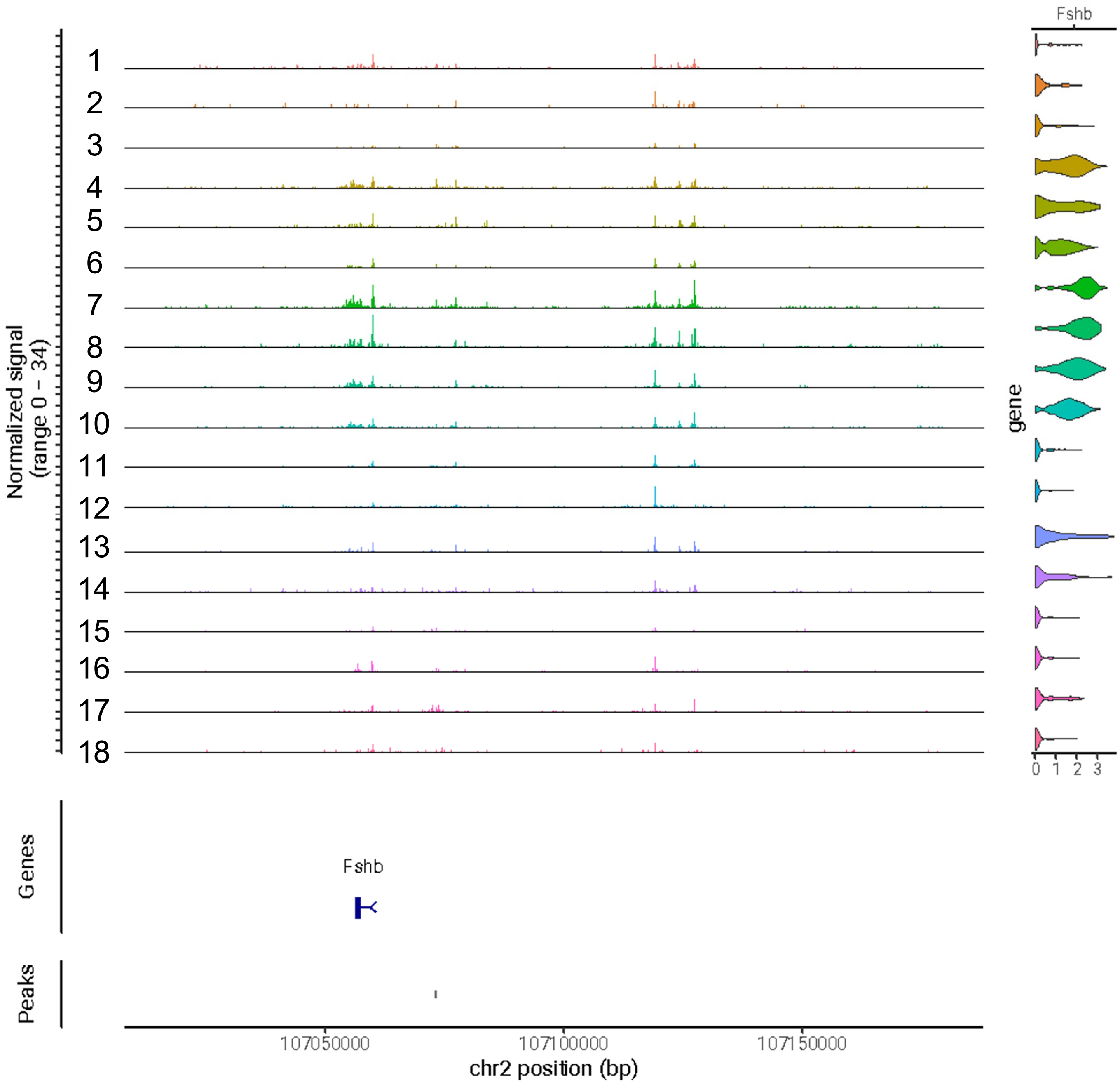

Fshb
Normalized signal
(range 0 - 34)
gene
0 1 2 3
Genes
Fshb
Peaks
107050000
107100000
107150000
chr2 position (bp)

Fig S10

**Figure S10. Chromatin accessibility upstream of the *Fshb* gene and *Fshb* mRNA expression across the different animals.** Shown are the chromatin accessibility tracks of the *Fshb* gene and 5' flanking region in each of the 18 animals (*Left*) and *Fshb* gene expression in the same samples (*Right*). Animal IDs are specified to the left of the figure. Presented at the bottom of the figure is a schematic view of the genomic region holding the *Fshb* gene and 5' flanking region, with genomic coordinates. Related to **Fig. 6**.

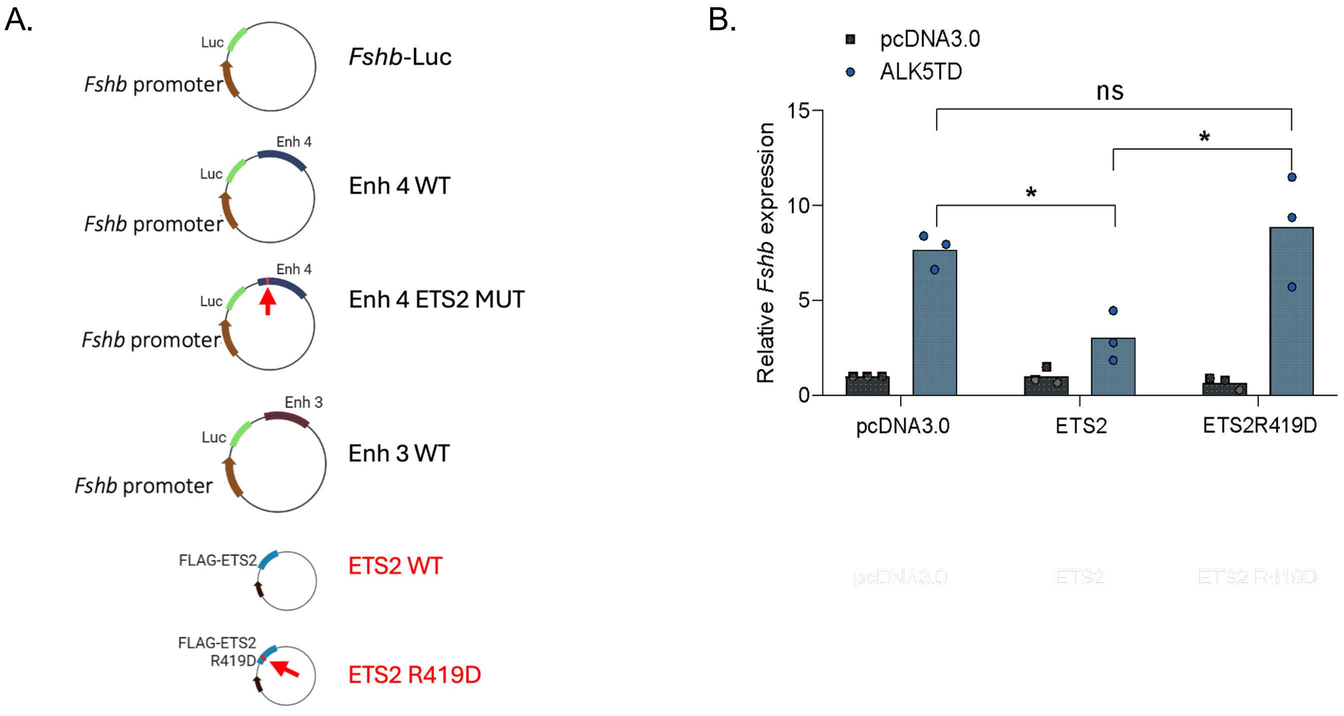
A.
Luc
Fshb promoter
Fshb-Luc
Enh 4
Luc
Fshb promoter
Enh 4 WT
Enh 4
Luc
Fshb promoter
Enh 4 ETS2 MUT
Enh 3
Luc
Fshb promoter
Enh 3 WT
FLAG-ETS2
ETS2 WT
FLAG-ETS2
R419D
ETS2 R419D
B.
pcDNA3.0
ALK5TD
ns
*
*
15
10
5
0
Relative Fshb expression
pcDNA3.0
ETS2
ETS2R419D

Fig S11

**Figure S11. Transfection constructs and additional experiments conducted in LβT2 cells.**
**A.** Schematic representation of the luciferase reporter constructs and the expression constructs used in transfection assays. Briefly, *Fshb*-luc contains the -1990/+1 mouse *Fshb* promoter region (brown arrow) inserted upstream of the luciferase gene (Luc; green segment); Enh4 WT and Enh3 WT have the Enh4 and the Enh3 region, respectively, inserted downstream of Luc, as depicted; in Enh4 Ets2 mut, the Enh4 region harbors an ETS2 binding site point mutation (see red arrow). Expression constructs are labeled in red and include: ETS2 WT, which encodes a FLAG-tag human ETS2 TF (blue segment); ETS2 R419D, encoding a FLAG-tag mutated human ETS2 TF (R419D mutation; see red arrow). **B.** LβT2 cells were co-transfected with 500 ng/well of either an empty expression construct (pcDNA3.0), ETS2 WT, or ETS2 R419D and 500 ng/well of either pcDNA3.0 or an HA-tag ALK5TD expression construct encoding a constitutively active ALK5 (ALK5TD). Endogenous *Fshb* gene expression was assessed by RT-qPCR. N = 3 independent experiments. Data were analyzed by Two-way ANOVA followed by Holm-Šidák test. *, $P<0.05$. Related to **Fig. 6**.

### Supplementary Tables

**Table S1.** Hormone levels and vaginal cytology-based staging of the estrous cycle

| Day of sacrifice | | | Animals | To detect LH surge<br>Whole Blood | | Terminal sample<br>Serum at sacrifice | | | | | | | | |
|---|---|---|---|---|---|---|---|---|---|---|---|---|---|---|
| Cohort 1 | Cohort 2 | | # | LH at 9 am | LH at 6 pm | LH (ng/ml) | FSH (ng/ml) | InhB (ng/ml) | InhA (ng/ml) | Age | | Strain | Vaginal cytology | Time of swab collection |
| 22-Jan | | 1 | 37 | 0.9722 | 22.8671 | 58.7146 | 3.3769 | 1.3958 | 1.5177 | 11 weeks | Mice acquired from commercial sources only inform the weeks | WT (C57BL/6) | Surging female at 6 pm (afternoon of proestrous) | 3 pm to 4 pm |
| 22-Jan | | 2 | 38 | 0.1164 | 19.0519 | 23.7029 | 1.7473 | 0.6512 | 1.9661 | 11 weeks | | WT (C57BL/6) | Surging female at 6 pm (afternoon of proestrous) | 3 pm to 4 pm |
| 22-Jan | | 3 | 39 | 0.2283 | 7.2562 | 16.6706 | 1.4592 | 1.2534 | 1.3469 | 11 weeks | | WT (C57BL/6) | Surging female at 6 pm (afternoon of proestrous) | 3 pm to 4 pm |
| | 17-Mar | 4 | 41 | 0.9427 | 15.8712 | 2.0587 | 2.2128 | 0.2135 | 0.0732 | 11 weeks | | WT (C57BL/6) | Surging female at 11 pm (afternoon of proestrous) | 3 pm to 4 pm |
| | 17-Mar | 5 | 42 | 0.3637 | 26.6473 | 4.4155 | 1.2563 | 0.2903 | 0.1574 | 11 weeks | | WT (C57BL/6) | Surging female at 11 pm (afternoon of proestrous) | 3 pm to 4 pm |
| | 17-Mar | 6 | 43 | 0.7231 | 22.4445 | 2.5412 | 1.4841 | 0.2672 | 0.0917 | 11 weeks | | WT (C57BL/6) | Surging female at 11 pm (afternoon of proestrous) | 3 pm to 4 pm |
| 20-Jan | | 7 | 44 | 0.2640 | 6.3431 | 5.4390 | 3.6371 | 0.3368 | 0.1090 | 11 weeks | | WT (C57BL/6) | Surging female at 2 am (early morning estrous) | 3 pm to 4 pm (day before) |
| 20-Jan | | 8 | 45 | 0.6299 | 29.5053 | 3.5018 | 3.0691 | not enough serum | | 11 weeks | | WT (C57BL/6) | Surging female at 2 am (early morning estrous) | 3 pm to 4 pm (day before) |
| 20-Jan | | 9 | 46 | 0.5334 | 8.9826 | 7.2728 | 2.4609 | 0.4371 | 0.1000 | 11 weeks | | WT (C57BL/6) | Surging female at 2 am (early morning estrous) | 3 pm to 4 pm (day before) |
| 20-Jan | | 10 | 47 | 0.7562 | 21.2741 | 6.5298 | 3.5889 | not enough serum | | 11 weeks | | WT (C57BL/6) | Surging female at 2 am (early morning estrous) | 3 pm to 4 pm (day before) |
| | 17-Mar | 11 | 50 | 1.3324 | 0.6788 | 5.1972 | 1.4269 | 1.1454 | 1.2551 | 11 weeks | | WT (C57BL/6) | Morning of proestrous, 9 am | within 1 hour before sacrifice |
| | 17-Mar | 12 | 51 | 2.0577 | 0.8990 | 5.1972 | 0.9628 | 1.2658 | 1.5734 | 11 weeks | | WT (C57BL/6) | Morning of proestrous, 9 am | within 1 hour before sacrifice |
| | 17-Mar | 13 | 54 | 2.1425 | ND | 4.5558 | 1.3316 | 1.2260 | 0.5690 | 11 weeks | | WT (C57BL/6) | Morning of proestrous, 9 am | within 1 hour before sacrifice |
| | 17-Mar | 14 | 55 | 0.3176 | 0.8990 | 4.8151 | 1.5243 | 0.9920 | 0.6614 | 11 weeks | | WT (C57BL/6) | Morning of diestrous, 9 am | within 1 hour before sacrifice |
| Excluded | 17-Mar | | 56 | 3.4087 | 1.7176 | 5.1011 | 1.4414 | 0.5072 | 0.1891 | 11 weeks | | WT (C57BL/6) | Morning of diestrous, 9 am | within 1 hour before sacrifice |
| | 17-Mar | 15 | 57 | 2.1848 | ND | 2.6265 | 0.6219 | 1.4037 | 0.6439 | 11 weeks | | WT (C57BL/6) | Morning of diestrous, 9 am | within 1 hour before sacrifice |
| | 17-Mar | 16 | 60 | ND | 0.2242 | 5.1732 | 1.7603 | 1.2335 | 0.3755 | 11 weeks | | WT (C57BL/6) | Morning of estrous, 9 am | within 1 hour before sacrifice |
| | 17-Mar | 17 | 61 | ND | 0.2242 | 6.7074 | 1.8067 | 1.5500 | 0.5299 | 11 weeks | | WT (C57BL/6) | Morning of estrous, 9 am | within 1 hour before sacrifice |
| | 17-Mar | 18 | 62 | 0.2242 | 0.5450 | 4.3689 | 1.7144 | 1.5827 | 0.3885 | 11 weeks | | WT (C57BL/6) | Morning of estrous, 9 am | within 1 hour before sacrifice |

Table S2: Samples and cells per estrous stage

| Stage | Proestrus_6pm | Proestrus_11pm | Estrus_2am | Proestrus_9am | Diestrus_9am | Metaestrus_9am | Estrus_9am | Total |
|---|---|---|---|---|---|---|---|---|
| Cell Count | 16379 | 19181 | 28230 | 33504 | 14416 | 15603 | 19477 | 146790 |
| Num of samples | 3 | 3 | 4 | 3 | 2 | 2 | 3 | 20 |

### **Table S3.** SnRNAseq (GEX) and snATACseq metrics

**GEX**

| Animal # | 1 | 2 | 3 | 4 | 5 | 6 | 7 | 8 | 9 | 10 | 11 | 12 | 13 | 14 | 15 | 16 | 17 | 18 |
|---|---|---|---|---|---|---|---|---|---|---|---|---|---|---|---|---|---|---|
| Aliquot_ID Animal | MF37 | MF38 | MF39 | MF41 | MF42 | MF43 | MF44_aggr | MF45 | MF46 | MF47 | MF50_aggr | MF51 | MF54 (multiplex) | MF55 | MF56 | MF60 | MF61 | MF62 |
| Estrous stage | Proestrus 6pm | Proestrus 6pm | Proestrus 6pm | Proestrus 11pm | Proestrus 11pm | Proestrus 11pm | Estrus 2am | Estrus 2am | Estrus 2am | Estrus 2am | Proestrus, 9 am | Proestrus, 9 am | Proestrus, 9 am | Diestrus, 9 am | Diestrus, 9 am | Estrus, 9 am | Estrus, 9 am | Estrus, 9 am |
| QC_result | Good | Good | Good | Good | Good | Good | Good | Good | Good | Good | Good | Good | Good | Good | Good | Good | Good | Good |
| GEX_QC_result | Good | Good | Good | Good | Good | Good | Good | Good | Good | Good | Good | Good | Good | Good | Good | Good | Good | Good |
| Not_Pass_metric | NA | NA | NA | NA | NA | NA | NA | NA | NA | NA | NA | NA | NA | NA | NA | NA | NA | NA |
| Estimated_number_of_cells | 9415 | 9512 | 20000 | 13415 | 9280 | 20000 | 20179 | 7773 | 11236 | 20000 | 27618 | 20000 | 16168 (2 samples) | 9095 | 7835 | 10134 | 15655 | 7264 |
| GEX_Mean_raw_reads_per_cell | 26343.101 | 28919.8294 | 31696.3514 | 17061.7805 | 55919.0224 | 28590.201 | 23775.49237 | 52116.565 | 52820.32 | 27601.2146 | 20437.40843 | 13188.1751 | 14375.058 | 23062.4065 | 24701.8521 | 28098.0001 | 14697.602 | 27068.6055 |
| GEX_Median_UMI_counts_per_cell | 4583 | 4455.5 | 3821 | 3195 | 5610 | 4550 | 4429 | 5661 | 5042.5 | 3727 | 4013.5 | 2380 | 2787 | 4053 | 3028 | 4397 | 2905 | 4621 |
| GEX_Reads_mapped_confidently_to_intronic_regions | 0.2809 | 0.3836 | 0.3269 | 0.279 | 0.3993 | 0.2826 | 0.263957026 | 0.3907 | 0.3283 | 0.3112 | 0.322065694 | 0.3602 | 0.2827 | 0.3223 | 0.3559 | 0.4565 | 0.3835 | 0.3379 |
| GEX_Reads_mapped_confidently_to_exonic_regions | 0.6092 | 0.5073 | 0.5536 | 0.6099 | 0.4879 | 0.582 | 0.628135589 | 0.4932 | 0.5266 | 0.5256 | 0.565585297 | 0.5158 | 0.5566 | 0.5623 | 0.5082 | 0.4312 | 0.4767 | 0.5457 |
| GEX_Q30_bases_in_read_2 | 0.942 | 0.9232 | 0.9489 | 0.943 | 0.9232 | 0.9488 | 0.938362667 | 0.9187 | 0.9473 | 0.9466 | 0.938803918 | 0.9389 | 0.9385 | 0.9403 | 0.941 | 0.9277 | 0.9335 | 0.9409 |
| GEX_Fraction_of_transcriptomic_reads_in_cells | 0.7182 | 0.794 | 0.6784 | 0.7044 | 0.786 | 0.6498 | 0.704029231 | 0.7987 | 0.6811 | 0.6777 | 0.749825503 | 0.7726 | 0.6725 | 0.7591 | 0.7345 | 0.847 | 0.7687 | 0.7435 |
| GEX_Over_20percent_mt_in_Cell | 0 | 0 | 0 | 0 | 0 | 0 | 0 | 0 | 0 | 0 | 0 | 0 | 0 | 0 | 0 | 0 | 0 | 0 |
| GEX_Percent_XIST | 0.9330 | 0.9449 | 0.9074 | 0.9433 | 0.9464 | 0.9597 | 0.9362 | 0.9541 | 0.9309 | 0.9241 | 0.9320 | 0.8310 | N/A | 0.9468 | 0.7832 | 0.9542 | 0.7011 | 0.9513 |
| GEX_Gender_determination | Female | Female | Female | Female | Female | Female | Female | Female | Female | Female | Female | Female | N/A | Female | Female | Female | Female | Female |

**ATAC**

| Animal # | 1 | 2 | 3 | 4 | 5 | 6 | 7 | 8 | 9 | 10 | 11 | 12 | 13 | 14 | 15 | 16 | 17 | 18 |
|---|---|---|---|---|---|---|---|---|---|---|---|---|---|---|---|---|---|---|
| Aliquot_ID Animal | MF37 | MF38 | MF39 | MF41 | MF42 | MF43 | MF44_aggr | MF45 | MF46 | MF47 | MF50_aggr | MF51 | MF54 (multiplex) | MF55 | MF56 | MF60 | MF61 | MF62 |
| Estrous stage | Proestrus 6pm | Proestrus 6pm | Proestrus 6pm | Proestrus 11pm | Proestrus 11pm | Proestrus 11pm | Estrus 2am | Estrus 2am | Estrus 2am | Estrus 2am | Proestrus, 9 am | Proestrus, 9 am | Proestrus, 9 am | Diestrus, 9 am | Diestrus, 9 am | Estrus, 9 am | Estrus, 9 am | Estrus, 9 am |
| ATAC_QC_result | Good | Good | Good | Good | Good | Good | Good | Good | Good | Good | Good | Good | Good | Good | Good | Good | Good | Good |
| ATAC_Q30_bases_in_read_1 | 0.922 | 0.943 | 0.936 | 0.937 | 0.937 | 0.936 | 0.940 | 0.942 | 0.928 | 0.930 | 0.941 | 0.937 | 0.9369 | 0.927 | 0.926 | 0.940 | 0.940 | 0.942 |
| ATAC_Q30_bases_in_read_2 | 0.925 | 0.925 | 0.930 | 0.922 | 0.927 | 0.932 | 0.925 | 0.928 | 0.939 | 0.935 | 0.923 | 0.918 | 0.9202 | 0.921 | 0.928 | 0.923 | 0.926 | 0.921 |
| ATAC_Confidently_mapped_read_pairs | 0.947 | 0.949 | 0.940 | 0.945 | 0.951 | 0.942 | 0.946 | 0.947 | 0.935 | 0.935 | 0.951 | 0.948 | 0.9473 | 0.948 | 0.943 | 0.953 | 0.948 | 0.945 |
| ATAC_Median_high.quality_fragments_per_cell | 15475 | 24404 | 10100 | 8620 | 14820 | 12960 | 15591 | 19102 | 15833 | 11389 | 11951 | 5681 | 8463 | 13279 | 16106 | 17489 | 7227 | 20170 |
| ATAC_Fraction_of_transposition_events_in_peaks_in_cells | 0.398 | 0.421 | 0.517 | 0.454 | 0.530 | 0.519 | 0.424 | 0.416 | 0.440 | 0.479 | 0.469 | 0.405 | 0.4753 | 0.401 | 0.456 | 0.438 | 0.427 | 0.416 |
| ATAC_TSS_enrichment_score | 9.307 | 9.292 | 12.003 | 10.400 | 12.260 | 12.301 | 9.391 | 10.026 | 10.710 | 11.591 | 9.680 | 9.889 | 11.0737 | 9.619 | 11.377 | 9.902 | 10.658 | 9.791 |

Table S4: Cell type proportions at each stage of the estrous cycle

| | Diestrus_9am | Proestrus_9am | Proestrus_6pm |
|---|---|---|---|
| **Corticotropes** | 0.0289 | 0.0370 | 0.0442 |
| **Gonadotropes** | 0.0189 | 0.0337 | 0.0237 |
| **Lacto-somatotropes** | 0.0000 | 0.0001 | 0.0000 |
| **Lactotropes** | 0.4616 | 0.4971 | 0.4699 |
| **Macrophages** | 0.0059 | 0.0076 | 0.0127 |
| **Melanotropes** | 0.0904 | 0.0704 | 0.0743 |
| **Pericytes** | 0.0141 | 0.0135 | 0.0131 |
| **Pit1-Lineage** | 0.0006 | 0.0005 | 0.0007 |
| **Pituicytes** | 0.0070 | 0.0047 | 0.0054 |
| **Proliferating** | 0.0031 | 0.0075 | 0.0039 |
| **Somatotropes** | 0.3267 | 0.2878 | 0.3005 |
| **Stem cells** | 0.0260 | 0.0174 | 0.0336 |
| **Thyrotropes** | 0.0170 | 0.0227 | 0.0180 |
